# Estimating Diversity Through Time using Molecular Phylogenies: Old and Species-Poor Frog Families are the Remnants of a Diverse Past

**DOI:** 10.1101/586420

**Authors:** O. Billaud, D. S. Moen, T. L. Parsons, H. Morion

## Abstract

Estimating how the number of species in a given group varied in the deep past is of key interest to evolutionary biologists. However, current phylogenetic approaches for obtaining such estimates have limitations, such as providing unrealistic diversity estimates at the origin of the group. Here we develop a robust probabilistic approach for estimating Diversity-Through-Time (DTT) curves and uncertainty around these estimates from phylogenetic data. We show with simulations that under various realistic scenarios of diversification, this approach performs better than previously proposed approaches. We also characterize the effect of tree size and undersampling on the performance of the approach. We apply our method to understand patterns of species diversity in anurans (frogs and toads). We find that Archaeobatrachia – a species-poor group of old frog clades often found in temperate regions – formerly had much higher diversity and net diversification rate, but the group declined in diversity as younger, nested clades diversified. This diversity decline seems to be linked to a decline in speciation rate rather than an increase in extinction rate. Our approach, implemented in the R package RPANDA, should be useful for evolutionary biologists interested in understanding how past diversity dynamics have shaped present-day diversity. It could also be useful in other contexts, such as for analyzing clade-clade competitive effects or the effect of species richness on phenotypic divergence. [phylogenetic comparative methods; birth-death models; diversity curves; diversification; extinction; anurans]

---

Estimating species diversity through geological time is key to our understanding of what controls biological diversity. Diversity curves have been extensively explored from fossil data and are at the origin of intensive debates on the role of stochasticity, diversity-dependence, and biotic and abiotic drivers on long-term diversity dynamics (Ezard et al., 2011; Ezard and Purvis, 2016; Foote et al., 2007; Liow et al., 2015; Marshall and Quental, 2016; Rabosky and Sorhannus, 2009; Silvestro et al., 2015).

Comparatively, only few studies have estimated analogous diversity curves from molecular phylogenies. Lineage-Through-Time (LTT) plots reporting the number of ancestral lineages in reconstructed phylogenies have been intensively used (see Ricklefs 2007 for a review), but these plots are missing all the lineages that did not leave any descendants in the present, thus giving the biased perception that diversity always increases steadily towards the present. While models of diversification that account for extinction started to be developed more than 25 years ago (Nee et al., 1992), these models and others with higher complexity have typically been used to estimate how speciation and extinction rates vary through time (see Morlon 2014; Pennell and Harmon 2013; Stadler 2013 for reviews), rather than to estimate diversity curves *per se.*

More accurately estimating diversity through time is important for understanding present-day patterns of species richness. One distinct pattern is that species richness can vary tremendously between closely related groups, but it is not clear why (Harmon, 2012). For example, the tuatara (Rhyn-cocephalia; *Sphenodon*) is a single extant species (Pough et al., 2015) whose sister group Squamata (snakes and lizards) has 10,417 species (Uetz et al., 2018). Part of this huge heterogeneity in species richness is almost certainly due to extinction in the tuataras, given an extensive fossil record (Jones et al., 2009) and hypotheses of competitive replacement by squamates (Apesteguía and Novas, 2003). By documenting differences in diversity over time, we can test such hypotheses about why one group is declining in diversity while the other is increasing, as well as identifying the time in the past at which the scale of diversity tipped from one group to another. Differences in extant diversity are seen across many groups such as whales (Morlon et al., 2011; Quental and Marshall, 2010), most salamander families vs. Plethodontidae, most snake clades vs. Colubroidea (Pough et al., 2015), which underlines the importance of understanding diversity curves through time, especially in groups with a poor fossil record.

Two of the first studies reporting species diversity through time with non-zero extinction estimated from molecular phylogenies were those of Morlon et al. (2011) and Etienne et al. (2011). These papers aimed to compare such curves to those estimated from the fossil record and to reconcile an apparent disagreement between paleontological and neontological estimates of diversity dynamics (Quental and Marshall, 2010). Morlon et al. (2011) obtained species-diversity curves by solving the deterministic differential equation that describes how the expected number of species varies with time under time-dependent diversification scenarios, using the maximum likelihood estimates of speciation and extinction rates (see Box 1 in Morlon 2014). This provided a first approach to estimating diversity curves, and hereafter we refer to it as the *deterministic* approach. However, this approach is approximate, it can lead to unrealistic diversity curves (some of which are illustrated in this manuscript), and it does not provide confidence intervals around diversity estimates. The approach proposed by Etienne et al. (2011) is similar, therefore sharing similar limitations. In addition, the latter approach assumes that diversification is diversity-dependent, such that species-diversity curves are constrained to increase and then reach a plateau over time, therefore excluding other types of dynamics such as those that include periods of diversity decline. Finally, while both approaches constrain current diversity and estimate backward in time, neither approach conditions its estimates on the known diversity at the root of the phylogeny, which must be one or two species (depending on whether the stem is included). Ignoring this conditioning can sometimes have dramatic effects on diversity estimates, as we will illustrate here.

In this paper we develop a more rigorous probabilistic approach for estimating Diversity-Through-Time (DTT) curves under time-dependent diversification models by deriving the full probability distribution of the number of species at each time point in the past. We test the performance of our new approach using intensive simulations. Finally, we apply our approach to three empirical cases: the cetaceans, which have become a model in the phylogenetic study of diversification (Condamine et al., 2013; Morlon et al., 2011; Quental and Marshall, 2010; Rabosky, 2014); Didelphidae (a family of American opossums), which yield an unrealistic diversity curve when using the deterministic approach; and anuran amphibians.

Anuran amphibians (frogs and toads; frogs here after for brevity) show stark diversity differences among clades. Frogs have been traditionally divided between the Archaeobatrachia (“archaic frogs”) and the Neobatrachia (“advanced frogs”, Duellman 1975; Ford and Cannatella 1993), with the latter nested within the former (Ford and Cannatella, 1993; Roelants and Bossuyt, 2005) and accounting for over 95% of all frog species (6721 of 7025 species; AmphibiaWeb 2016; Pough et al. 2015). Individual families within these groups show similar patterns: eight of the 10 archaeobatrachian families have less than 12 species, whereas 34 of the 44 neobatrachian families each have higher diversity than this and 12 families have over 200 species each (AmphibiaWeb, 2016). Furthermore, most older neobatrachian families have very low diversity (with ‘older” referring to the stem age of the family, Feng et al. (2017); Pyron (2014); Pyron and Wiens (2011)). Finally, the scarcity of the anuran fossil record means that paleontological methods for assessing these diversity differences through time are not possible. Thus, frogs are an excellent group to examine the utility of our approach. Note that while Archaeobatrachia is paraphyletic (cRoelants and Bossuyt, 2005), we use it here as a concise term that represents an informal group of old anuran families (Ford and Cannatella, 1993).

## METHODS

### Probability distribution of the number of species in the past

We assume that a clade comprising *n* species at present has evolved from a single lineage according to a birth-death model of cladogenesis (Nee et al., 1992), with per-lineage speciation and extinction rates, *λ*(*t*) and *μ*(*t*), respectively, that can vary over time. We note *N* (*t*) the number of species at time *t*, with *t* measured from the past to the present (the time at present is denoted *T*_max_, and corresponds either to the crown or stem age; thus *N*(*T*_max_) = *n*). We consider the phylogeny of *l* species sampled at present from this clade, which can be fewer species than the entire clade (i.e. *l < n*).

If we have *a priori* knowledge of the total number of species in the clade *n*, we can compute the probability that there were *m* species at a given time *t*, knowing that there are *n* extant species today, and that there were *x* species at an earlier time *s* (*s* < *t*). We show (Appendix) that this probability is given by:

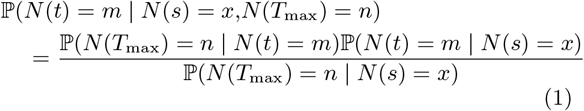

with

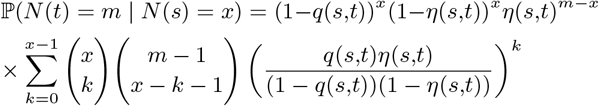

where *q*(*s,t*) is the probability that a lineage alive at time *s* goes extinct between *s* and *t* and *η*(*s, t*) is the probability that a lineage alive at time *s* gives birth to two lines that survive to time *t*. These latter probabilities are given by Kendall (1948):

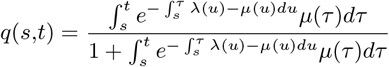

and

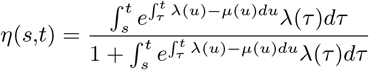

We also provide a second formula corresponding to a hypothetical case when we have information on the fraction *f* that an extant species has been sampled (*f* < 1) rather than on the total number of species *n*. While this is less common, we anticipate that likelihood methods for studying diversification when the total number of species is unknown will soon be developed (Lambert, 2018), in particular in order to study the diversification of microbial groups (Lewitus et al., 2018; Louca et al., 2018; Morlon et al., 2012). Such approaches will directly estimate the fraction of species sampled rather than the total number of species, and in this case it will be more accurate to use this direct estimate. We can compute the probability that there were m species at a given time *t*, knowing that there are *l* extant species represented in the phylogeny, and that there were *x* species at time *s* (*s* < *t*). We show (Appendix) that this probability is given by:

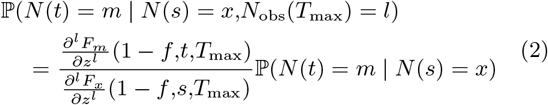

where *F_x_*(*z, s, t*) is the probability-generating function for 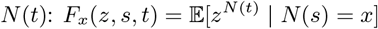, and its derivatives are given by:

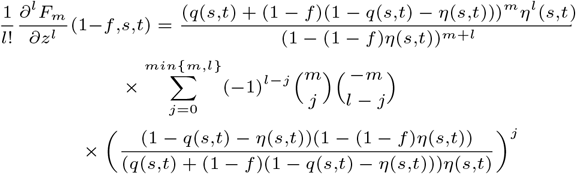

with *q*(*s,t*) and *η*(*s,t*) as above.

Equations (1) and (2) both provide us with an analytical formula for the probability distribution of the number of species in the past. While these expressions are valid for all *s* < *t*, we use them here to force the number of species at the origin of the clade to be 1 (if *T*_max_ is the stem age) or 2 (if *T*_max_ is the root age) and thus fix *s* = 0 and *x* is either 1 or 2. Missing species in the phylogeny do not affect the stem age of the clade, but they might affect its crown age. We assume here for simplification that the crown age is not greatly affected by undersampling, which is likely to be the case for moderate levels of undersampling (Sanderson, 1996). In what follows, we focus on the case when there is knowledge on the total number of species in the clade n, and we use equation (1).

Given the phylogeny of *l* species sampled at present, and under the hypothesis that diversification rates are identical across lineages, the probability distribution of the number of species in the past is obtained in two steps. First, we need to estimate how the speciation and extinction rates (λ(*t*) and μ(*t*), respectively) vary through time. Next, we compute 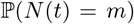 for a pre-defined series of times *t* and for each *m* value. In the first step, we estimate λ(*t*) and μ(*t*) by maximum likelihood, finding both the functional form (e.g. constant, linear, exponential) of the time-dependency of the rates and the associated parameters that maximize the likelihood given the phylogeny (Morlon et al., 2011). We perform these analyses using the *fit_bd* function from the R package RPANDA (Morlon et al., 2016). These return estimates of λ(*t*) and *μ*(*t*) with *t* measured from the present to the past. In the second step, we compute the probability associated with each *t* and *m* using the formulas above. From those, we obtain for each time *t*: *i*) the expected number of species by computing 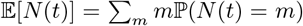 and *ii*) the confidence interval around this expected value by keeping the values of *m* with highest probability that collectively sum up to 0.95 and discarding the remaining *m* values. Codes for these analyses are freely available on GitHub (https://github.com/hmorlon/PANDA) and included in the R package RPANDA (Morlon et al., 2016). The *prob_dtt* function computes the probabilities, and the *plot_prob_dtt* function computes and plots the expected values and confidence intervals around them. The *m* values for which a probability is computed at each time *t* are chosen by the user of the function, and will typically be all integers from 1 to *m_max_*, with *m_max_* such that the sum of probabilities is almost equal to 1. Here, we chose the *m_max_* values such that the sum of probabilities is at least 0.99.

In our Appendix, we provide an analytical solution for computing the expected number of species under the birth-death process conditioned on the number of species at the root, and an alternative procedure for obtaining confidence intervals. We did not use these results here, as it was computationally more efficient to use the already computed probability distribution. We also provide analytical solutions for the rates of the conditioned birth-death process; these solutions could for example be useful for efficiently simulating specific realizations of DTT curves.

The simple models considered above may be poor approximations of the real diversification process occurring in nature, in particular for old clades that are ecologically and phenotypically diverse and that have experienced major extinction events and/or dramatic environmental changes. There are many ways that diversification processes can deviate from these simple models. A common feature of diversification rates is to vary across lineages, and in this case applying homogeneous birth-death processes can lead to spurious inferences of past dynamics (Morlon et al., 2011; Rabosky, 2010). Extending our approach in order to account for known rate heterogeneities is straightforward: following what was done in (Morlon et al., 2011), one can analyze clades with different diversification regimes separately, compute each of their DTT curves, and sum them up to obtain a global DTT curve. While detecting shifts in diversification regimes without any a priori hypothesis on where the shifts might occur is challenging (Alfaro et al., 2009; Moore et al., 2016; Rabosky, 2014), testing for the presence of shifts at specific locations in the phylogenies, such as at the origin of specific clades, can be done using classical model selection (Morlon et al., 2011). Specific models have been developed to account for other types of deviations, such as mass extinction events (Stadler, 2011) and environmental changes (Condamine et al., 2013; Lewitus and Morlon, 2017). Once estimates of diversification rates through time are inferred from such models, they can be used in our equations to obtain DTT curves, although we have not implemented this here.

### Testing the performance of the approach

We thoroughly tested the performance of our approach using simulations, starting with the case of homogeneous diversification dynamics. We simulated three types of diversity curves corresponding to expanding diversity (species richness increases towards the present), waxing-waning diversity (species richness increases and then decreases towards the present), and saturating diversity (species richness increases before oscillating around an equilibrium value). The expanding scenario was simulated with constant speciation and extinction rates. The waxing-waning and saturating scenarios were both simulated with either exponentially decreasing speciation towards the present and constant extinction, or constant speciation and exponentially increasing extinction towards the present, producing a total of five simulation scenarios (Fig. 1). Parameter values used in the simulations were randomly drawn from a uniform distribution for each simulation. We fixed the simulation time to 150 Myr. In order to obtain trees of realistic and manageable size under each scenario, we used the following constraints (here t runs forward, from the past to the present; *T_max_* = 150):

- expanding: λ ∈ [0.05,0.1] and *μ* ∈ [0,λ].
- waxing-waning (speciation decreasing): λ(*t*) = *a* exp(−*bt*) with *a* ∈ [0.1,0.2], *μ* ∈ [0,*a*/2], and b such that *t*_eq_ satisfying λ(*t_eq_*) = *μ* is in 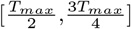.
- waxing-waning (extinction increasing): λ ∈ [0.05,0.2], *μ*(*t*) = *a* exp(*bt*) with *a* ∈ [0,λ] and b such that *t_eq_* satisfying *μ*(*t_eq_*) = λ is in 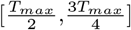. *μ*(*T_max_*) ≥ λ.
- saturating (speciation decreasing): λ(*t*) = *a* exp(−*bt*) + *μ* with *μ* ∈ [0,0.5], *a* ∈ [0,2*μ*] and b such that *t_eq_* satisfying exp(-*bt_eq_*) = 0.001 is in 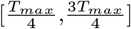.
- saturating (extinction increasing): λ ∈ [0.05,0.1], *μ*(*t*) = *a* exp(*bt*) with 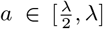 and b such that *μ*(*T_max_*)=λ.

**Figure 1:**
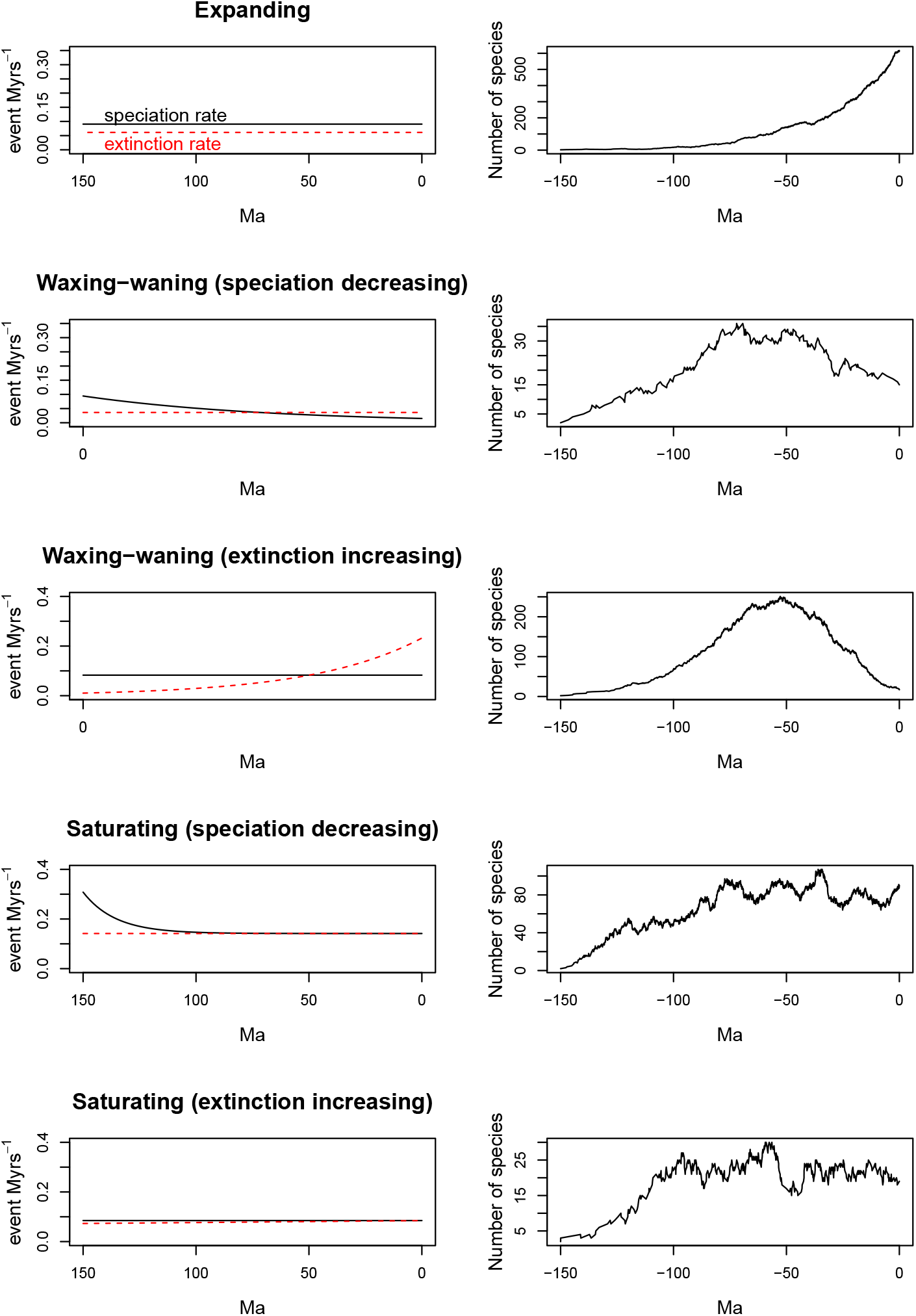
Diversification scenarios used in our simulations and corresponding diversity dynamics. Left panels: a specific realisation of rates of speciation (black lines) and extinction (red dashed lines) through time used in our simulations; right panels: a specific realisation of diversity through time under each diversification scenario, simulated with the parameters shown on the left.

Figure 1 illustrates one realisation of each scenario. We used the simulation approach of Paradis (2011) implemented in the *rlineage* function of the R package APE to obtain complete phylogenies (with extinct species) and the *ltt.plot.coords* function to obtain the reconstructed phylogenies (without extinct species). We discarded trees with less than 10 or more than 10,000 tips (this resulted in a less than 10% rejection rate) and simulated 400 phylogenies for each of the five scenarios. For each phylogeny and each 1 Myr time step (between 0 and 150), we recorded the observed (simulated) number of extant species and three different estimates of species diversity: *i*) our new probabilistic approach described above, *ii*) the deterministic approach of Morlon et al. (2011), and *iii*) the number of lineages on the reconstructed phylogeny (i.e. the well-known lineage-through-time plot, an estimate that ignores extinctions). For *i*) and *ii*), we first selected the model, among the 5 described above, that gave the best support (i.e. had the lowest AIC score) given the data (the model providing the best support was not necessarily the generating model) before computing the corresponding DTT curve. We used the crown age condition for both the deterministic and probabilistic approach. Finally, we measured a global error 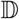 between the observed (denoted *obs*) and estimated (or theoretical, denoted *th*) diversity curves by averaging the relative error over time:

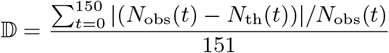

Fig. S1 illustrates examples of simulated diversity-trajectories and estimated DTT curves (with confidence intervals), along with associated global errors, under each of the five diversification scenarios.

We also separated the error corresponding to an overestimation of the number of extant species from the error corresponding to under-estimation. We did this by counting both the number of overestimates and underestimates along the diversity curve and the magnitude of each type of error. The magnitude of the overestimation was measured as

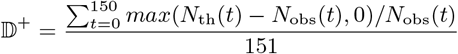

And that of the underestimation as

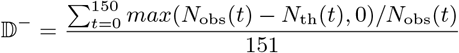

such that 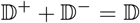.

We analyzed the effect of undersampling (missing species in the phylogeny) on diversity estimates. We pruned the simulated phylogenies described above to a fraction of 0.75, 0.5 and 0.25, estimated DTT curves for each phylogeny and each sampling fraction, and computed the resulting global error. For comparison, we also computed this error for the two other analytical approaches (i.e. the LTT and the deterministic approach). Here again we used the crown age condition; undersampling may lead to an underestimation of the crown age and it is valuable to evaluate the potential bias introduced by this underestimation given that empirical analyses often ignore the effect of undersampling on crown age estimates. We also analyzed the effect of tree size on the ability to properly estimate species richness by reporting 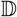 computed on complete trees binned on a log_2_ scale according to their size.

As discussed earlier, there are many ways in which diversification processes can deviate from the simple models tested here, and we cannot thoroughly assess the effect of all of them. We analyzed the performance of the method when diversification is not homogeneous across lineages. We simulated > 400 trees under an expanding scenario (constant speciation and extinction rates) but with a shift in diversification rates happening 50 Ma. We performed these simulations with our own codes by simulating a 150 Myrs old phylogeny with diversification rates randomly chosen as above, selecting the node the closest to −50 Myrs, and replacing the clade descending from this node by a phylogeny of the corresponding age simulated with a new set of randomly chosen diversification rates. For each phylogeny we tested whether there was a significant support for the shift following the approach of (Morlon et al., 2011), and computed resulting diversity curves (i.e. a single DTT curve if no shift was detected, and the sum of two independent DTT curves if a shift was detected) and global errors. We also explored the bias that might occur by artifactually detecting inexistent shifts: we simulated > 400 expanding trees (same parameters as above) with no shift, tested support for a shift (we performed this test at the node just following 50 Ma that subtended the most species and thus is the most likely to support an non-existant shift), and computed resulting diversity curves and global errors as above.

Finally, we analyzed how well the method performs when events occur that are not accounted for by our model, such as mass extinction events. We simulated > 400 150 Myrs old trees under an expanding scenario with a mass extinction event happening 50 Myrs ago, using the *sim.rateshift.taxa* function of the TreeSim R package (Stadler, 2015). The background diversification rates were sampled as above and the proportion of species surviving the mass extinction event was uniformly sampled in [0.1,0.9]. For each phylogeny we computed DTT plots and global errors (without testing for the presence of potential shifts).

### Empirical Applications

In order to illustrate the utility of our approach and to compare it to the deterministic one, we considered three empirical applications. First, we analyzed diversity curves inferred from the cetacean phylogeny (Steeman et al., 2009); diversity-through-time curves for this group have been estimated in both Morlon et al. (2011) (see their Figure 1a) and Etienne et al. (2011). Morlon et al. (2011) showed that diversification dynamics were not homogeneous across cetaceans, and in particular that the four most species-rich cetacean families (Balaenopteridae, Delphinidae, Phocoenidae and Ziphiidae) and the ‘‘backbone”, defined here as the phylogeny composed of the other cetacean species, diversified with distinct models and rates. Hence, following Morlon et al. (2011), we computed separate diversity curves for these distinct parts of the tree. The cetacean phylogeny is missing one species from Delphinidae and one from Ziphiidae. We accounted for these missing species when estimating λ(*t*) and μ(*t*) for these groups. We used the stem age condition (*n* = 1) for the four families and the crown age condition (*n* = 2) for the remaining cetaceans (we did not have information about the stem age in the cetacean phylogeny). Second, we analyzed the phylogeny of Didelphidae, a family of American opossums comprising 100 extant species (74 of which are represented in the phylogeny, *f* = 0.74), which yields an unrealistic diversity curve when using the deterministic approach (see Results). We took this phylogeny from the updated version of the mammalian trees of Faurby and Svenning (2015) (66 of the 74 species in the tree have molecular data). For these two empirical examples, we computed diversity-through-time curves using the deterministic approach of Morlon et al. (2011), as well as probability distributions, expected diversity-through-time curves, and confidence intervals around these curves using the new probabilistic approach.

Finally, we wanted to examine the utility of our method for understanding patterns of diversity through time in groups that may have low diversity due to extinction, but for which there are few fossil data. For this we estimated the diversity dynamics of frog families in Archaeobatrachia, many of which are older but show lower diversity than the families of the more recent Neobatrachia. If diversity dynamics were homogeneous over anuran history, then these older groups would have higher diversity than the more recent groups. We estimated diversification rates for archaeobatrachian families and reconstructed their history of diversity over time, considering the possibility of extinction. Estimating extinction rates with extant clades (i.e. without fossils) is contentious (Rabosky, 2010), yet key studies have found that one can reasonably estimate extinction given appropriate methods (Morlon et al., 2011) and conditions (Beaulieu and O’Meara, 2015).

We focused on the phylogeny of Archaeobatrachia and used the amphibian phylogeny from Pyron (2014), which contains 135 of the 264 species (*f* = 0.51) from this group and was the most completely sampled time-calibrated anuran phylogeny available at the time of our analyses. Jetz and Pyron (2018) recently published a tree with nearly all described anuran species. We expect that analysing this phylogeny would produce similar results, given that (1) most additional species in the fully sampled tree were semi-randomly imputed (based on taxonomy) onto a smaller molecular phylogeny; (2) the molecular datasets and phylogeny estimation methods of the two papers (Jetz and Pyron, 2018; Pyron, 2014) are highly overlapping, thus likely producing very similar molecular-data-only trees; and (3) Jetz and Pyron (2018)‘s diversification analyses of the fully sampled tree gave similar results as their analyses based on the tree based on molecular data alone.

Given that extinction can be masked by a stronger statistical signal from recently radiating clades (Morlon et al., 2011), we assumed that there could be shifts in diversification rates, and that these shifts occurred at the base of families (i.e. the beginning of their stem branches). There are 10 families in Archaeobatrachia, nine of which have two or more species. We considered the possibility of a maximum of nine rates shifts, each happening at the base of one of these families. We followed a stepwise procedure of shift selection, meaning that we first tested statistical support for a single rate shift producing two distinct diversification regimes within Archaeobatrachia, each with its own best-fit diversification model. Here we tested constant rates and rates that varied as an exponential function of time. If there was support for a single rate shift, we assigned this shift to the family that showed the highest improvement in the overall likelihood. We iterated the process to examine whether sufficient statistical support existed for additional rate shifts, until there was no statistical support for further partitioning the overall model of diversification. At each step, statistical support was assessed using a likelihood ratio test. Finally, we estimated diversity trajectories for each group that had independent diversification dynamics, using our new probabilistic approach. We used clade-specific sampling fractions: *f* = 1 for Ascaphidae, Bombinatoridae, Leiopelmatidae, Pelobatidae, Pelodytidae and Scaphiopodidae, *f* = 0.92 for Alytidae, *f* = 0.73 for Pipidae, and *f* = 0. 36 for Megophryidae. The backbone phylogeny that subtended the five families with rate shifts had a sampling fraction *f* = 0.96. Each sampling fraction was computed as the number of species represented in the phylogeny divided by the known number of species in the clade.

## 1 RESULTS

### 1.1 Performance of the approach

Our simulation analyses showed that the new probabilistic approach improves the accuracy of diversity estimates (Fig. 2). As expected, the LTT plot, which ignores extinct lineages, performs the worst on average. The deterministic approach tends to improve diversity estimates, but not always. The probabilistic approach outperforms all previous approaches, with a reduced global error for all diversification scenarios. The improvement is the most notable in the waxing-waning and saturating scenarios. All three methods tend to underestimate rather than overestimate species diversity through time in these curves (Fig. S2 & S3). The probabilistic approach was more robust to undersampling than either the LTT or the deterministic approach, under all diversification scenarios (Fig. 2). There was no clear effect of tree size on the global error 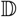 obtained with the probabilistic approach (Fig. 3). The approach performed as well in the presence of a shift in diversification rates, with a median error of 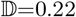, and when there was no shift but we tested for the presence of one (median 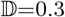). The error increased when mass-extinction events were simulated but unmodeled, but only slightly so (median 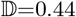).

**Figure 2:**
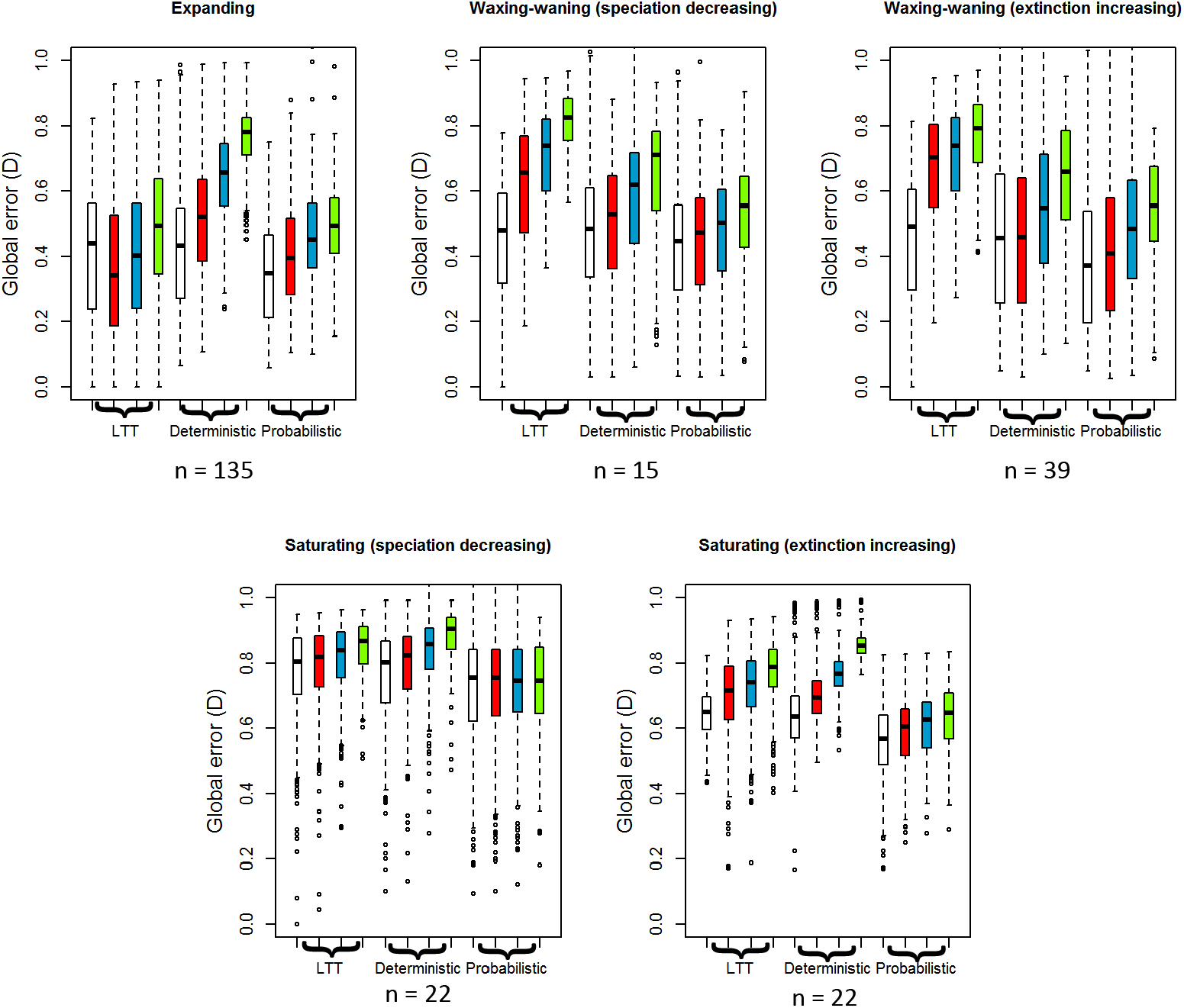
Accuracy of diversity-through-time estimates and effect of undersampling. Global error 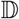 for trees simulated under the five diversification scenarios considered in the paper when using each of the three diversity-through-time estimates: the Lineage-Through-Time (LTT) plot, the deterministic estimate, and the expected value of diversity provided by the probabilistic approach. Inference for complete trees are represented in white and colors represent the degree of undersampling (red: sampling fraction of 75%, blue: 50%, green: 25%). Boxplots represent the median, 1^*st*^ and 4^*th*^ quartile over 400 simulations, whiskers represent the lowest (and highest) datum still within 1.5 interquantile range of the lower (resp.upper) quartile, and dots represent outliers. n values are the median number of extant species in the complete trees.

**Figure 3:**
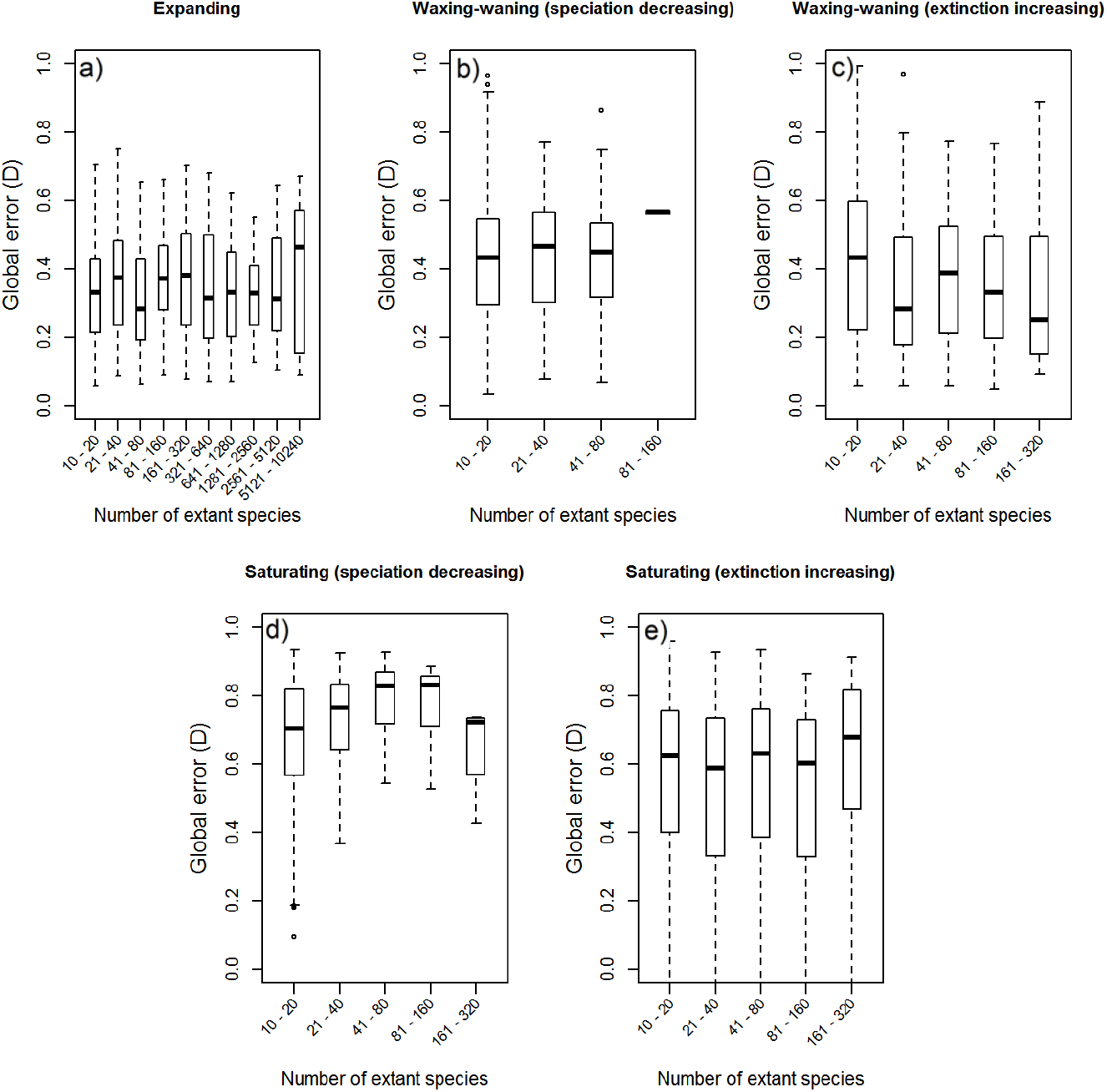
Effect of tree size on the accuracy of diversity-through-time estimates. Global error 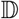 for trees simulated under the five diversification scenarios considered in the paper, binned according to their size, and for diversity estimates computed as the expected value of diversity provided by the probabilistic approach. Boxplots represent the median, 1^*st*^ and 4^*th*^ quartile over 400 simulations, whiskers represent the lowest (and highest) datum still within 1.5 interquantile range of the lower (resp.upper) quartile, and dots represent outliers.

### 1.2 Cetacea and Didelphidae

Our new approach for computing the expected number of species recovered DTT curves for the backbone cetacean phylogeny and for the four richest cetacean families that closely matched the ones obtained with the deterministic approach of Morlon et al. (2011) (Fig. 4b); the resulting diversity dynamics for the cetaceans were shown in the latter paper as consistent with fossil data. Consistent with Morlon et al. (2011), we found that the best-fit model for the backbone cetacean phylogeny was a model with constant speciation rate (λ estimated at 0.23) and increasing extinction rates towards the present (*μ*(*t*) = 0.9*e*^−0.15(*T_max_−t*)^); this results in a diversity-through-time curve that increases until ~ 10 Ma, reaches a diversity peak at this time, and then declines. In addition, our new approach provides a confidence interval around the diversity-through-time curve that shows that even the lower bound of the diversity curve supports a waxing-waning diversity pattern with a peak of cetacean diversity ~ 10 Ma (Fig. 4a).

**Figure 4:**
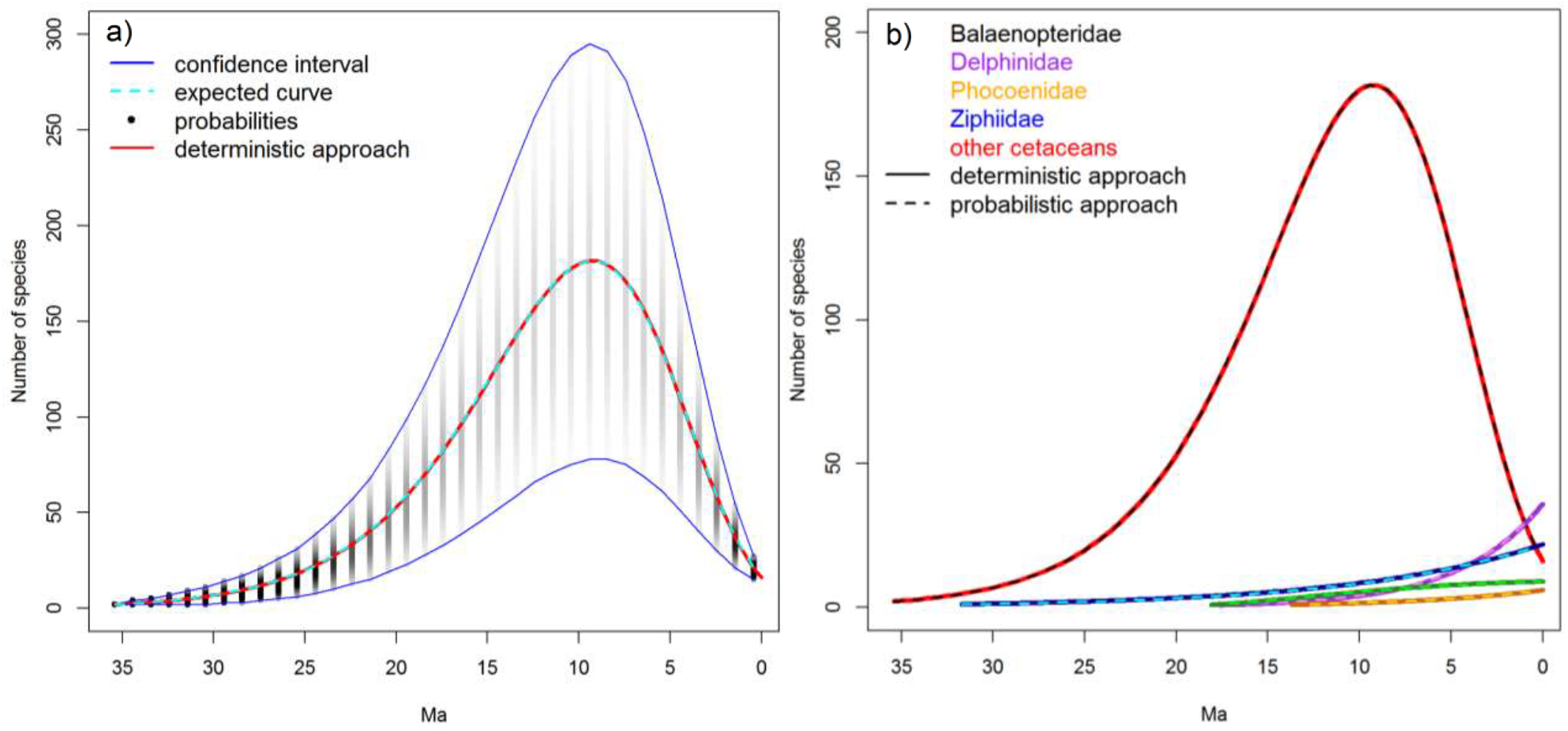
Estimated diversity-through-time curves for the cetaceans. a) Probability distribution (in black, the color intensity reflects probability values), expected value (in cyan, dashed curve), and confidence interval (in blue) of the number of extant species at each 1 Myr interval for the *backbone* cetacean phylogeny (i.e. a phylogeny that excludes the four main cetacean families); the diversity-through-time curve provided by the deterministic approach is plotted for comparison (solid red curve) b) Comparison between diversity curves obtained with the deterministic approach (solid curves) and the expected diversity-curves obtained with the probabilistic approach (dashed curves) for the four main cetacean families and the backbone phylogeny (referred to as *other cetaceans*).

The best-fit model for the Didelphidae was a model with constant speciation rate (λ estimated at 0.12) and decreasing extinction rate towards the present (*μ*(*t*) = 0.0041*e*^0.091(*T_max_−t*)^). With these estimates of diversification rates, the deterministic approach infers an unrealistic diversity-through-time curve with a ridiculously high number of species at the origin of the group(Fig. 5a). This is a situation that we encountered on several occasions in other analyses, and it likely comes from the amplified effect of (even small) biases in diversification rate estimates on diversity curves when species richness at the origin of the group is not constrained. In contrast, our new probabilistic approach that constrains species richness at the origin of the group provides realistic estimates, as illustrated here for Didelphidae (Fig. 5b).

**Figure 5:**
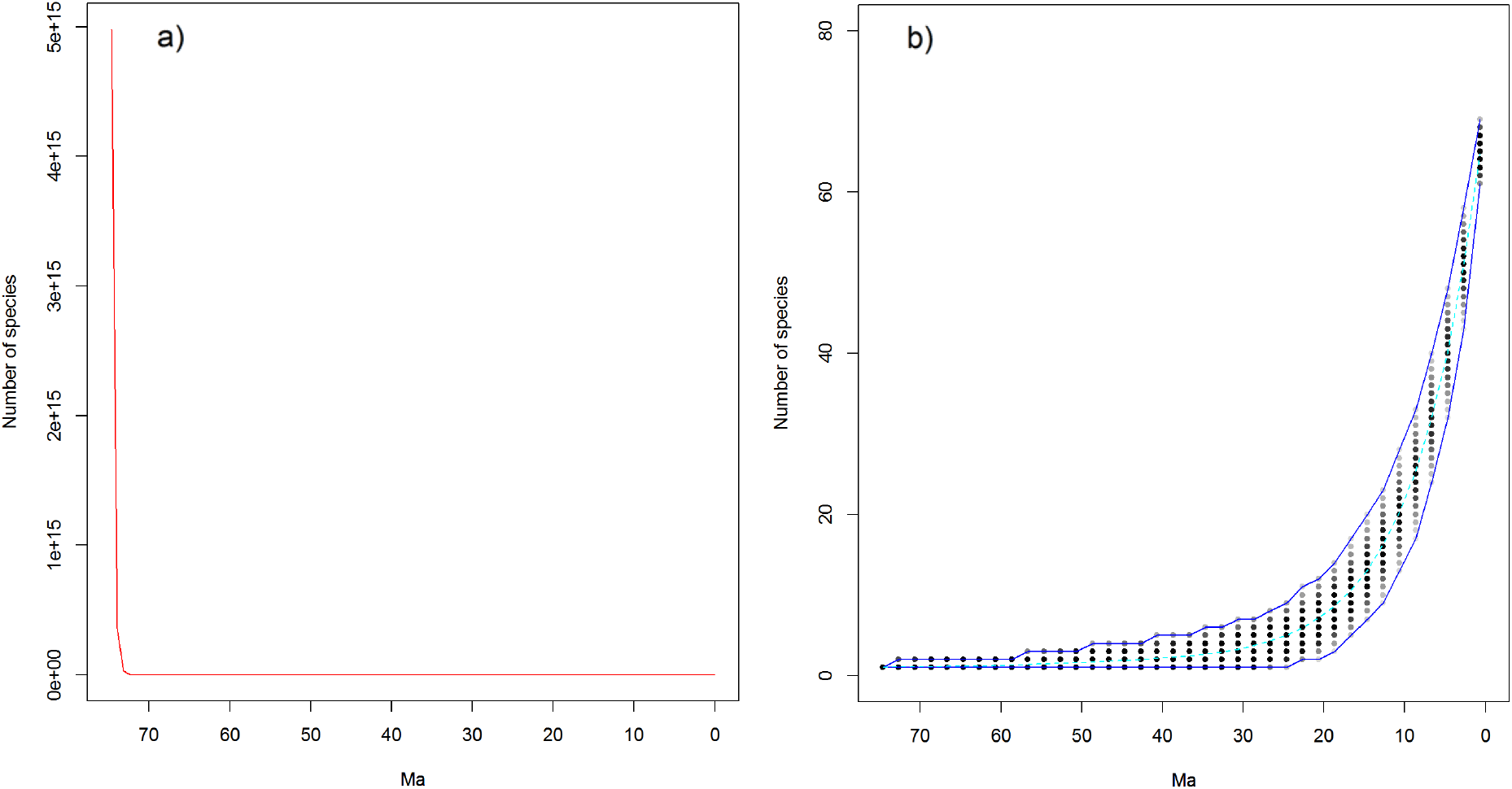
Estimated diversity-through-time curves for Didelphidae. a) Deterministic diversity-through-time curve b) Probability distribution (in black, the color intensity reflects probability values), expected value (in cyan, dashed curve), and confidence interval (in blue) of the number of extant species at each 1 Myr interval for the Didelphidae.

### 1.3 Diversity through time of frogs

We found evidence for five shifts in diversification dynamics in Archaeobatrachia at the base of Megophryidae, Bombinatoridae, Pelodytidae, Pipidae and Pelobatidae (Table S1). Past diversity of Archaeobatrachia was much higher than current diversity, reaching a peak of diversity up to ~ 2530 species around 166 million years ago (Fig. 6). The wax-wane pattern of diversity observed in the backbone phylogeny (which includes Alytidae, Scaphiopodidae, Leiopelmatidae, Ascaphidae, and Rhinophrynidae) was robust even if we discarded some of the inferred diversification rate shifts, though the exact diversity estimates depended on how many diversification shifts were assumed (Fig. S4). The diversity decline in the backbone phylogeny was due to both speciation and extinction rates declining over time, with a faster slowdown in speciation than extinction (Fig. S5). The five families subtending rate shifts were all expanding in diversity, but with distinct diversification scenarios (Fig. S5): Megophryidae, Bombinatoridae and Pipidae experienced very little extinction, while Pelodytidae and Pelobatidae had high extinction rates at the beginning of their histories that resulted in long stem branches.

**Figure 6:**
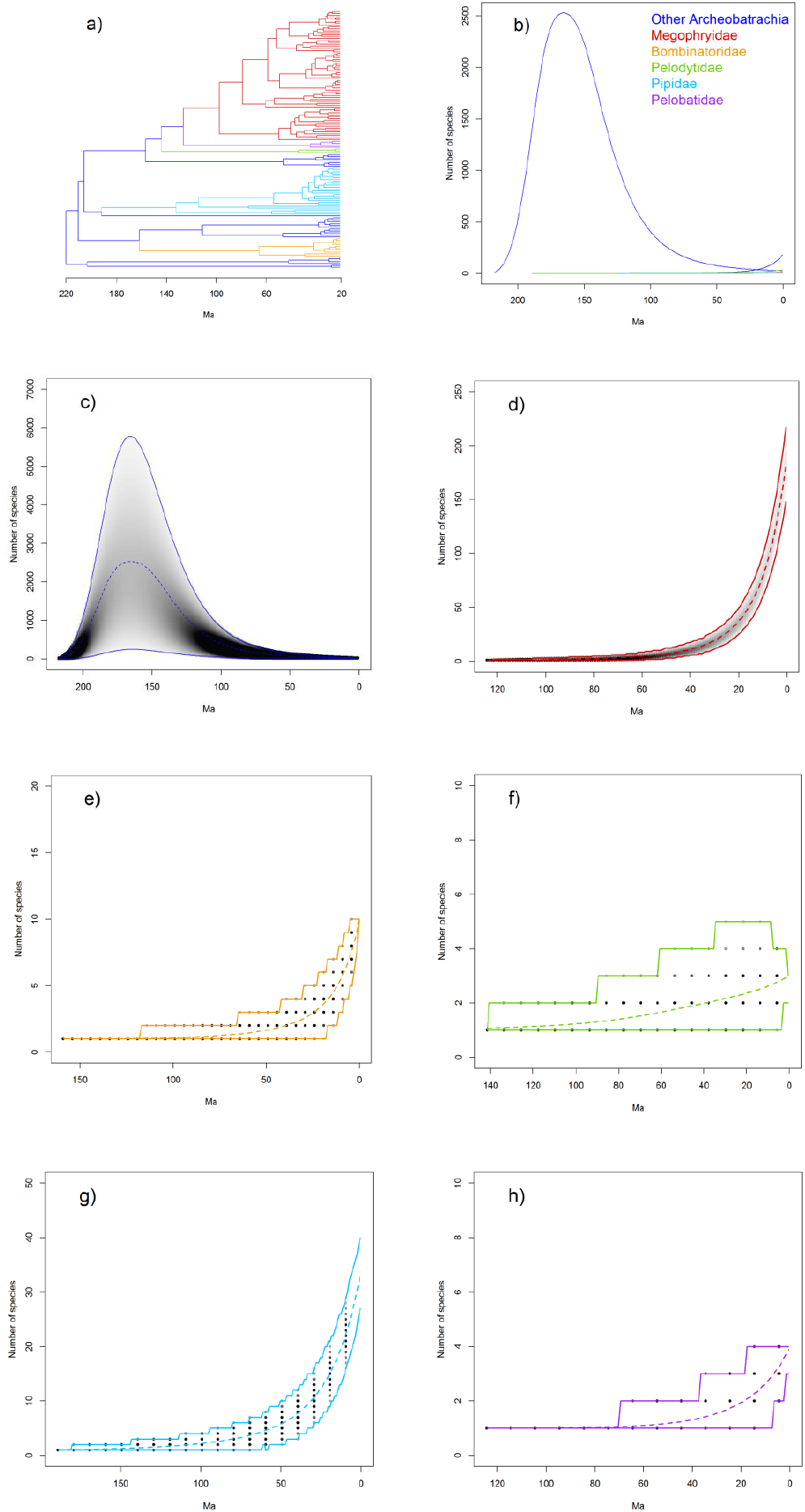
Estimated diversity-through-time curves for Archaeobatrachia. a) The phylogeny of Archaeobatrachia b) Estimated diversity through-time (expected value) for the *backbone* Archaeobatrachia phylogeny (i.e. a phylogeny that excludes the five archaeobatrachian families subtending diversification rate shifts, referred to as *Other Archaeobatrachia*) and the five archaeobatrachia families subtending diversification rate shifts. c-h) Probability distribution, expected value, and confidence interval of the number of extant species at various time points for the *backbone* archaeobatrachia phylogeny and the five archaeobatrachian families subtending diversification rate shifts. The probability distribution, expected value, and confidence interval is plotted at each 1 Myr interval for the backbone and the Megophryidae, but at larger intervals for the other groups for better presentation purposes.

## DISCUSSION

We derived probability distributions for the number of extant species in the past. Given the phylogeny of a group, these expressions provide estimates of how the species richness of this group varied through time and a confidence interval around these estimates. We implemented these expressions in the R-package RPANDA (Morlon et al., 2016), which should help evolutionary biologists derive diversity curves for groups of interest.

We provided (and implemented) two expressions, the first one corresponding to the case when there is *a priori* knowledge of the total number of extant species in the clade, and the second one corresponding to the case when there is *a priori* knowledge of the probability that an extant species is represented in the phylogeny. In practice, current likelihood models of diversification require providing a sampling faction, which is computed by dividing the number of species represented in the phylogeny by the total number of species known for the group. When this total number of species is known (or rather well estimated, which is often the case for macroorganisms), the first expression should be used, even if some species are not represented in the phylogeny. However, there are cases, in particular when studying microorganisms, when obtaining robust estimates of total diversity is challenging. So far, the few studies applying diversification models to microbial groups have either assumed a very wide range of total diversity values (Morlon et al., 2012), or used estimates of total diversity obtained from mark-recapture-type techniques (Louca et al., 2018) or Bayesian extrapolations of rank abundance curves (Lewitus et al., 2018; Quince et al., 2008). In the future, we anticipate that likelihood methods for studying diversification when the total number of species is unknown will be developed (Lambert, 2018), in particular to deal with such microbial groups. Such approaches will not require *a priori* knowledge of the total diversity and will directly estimate the fraction of species sampled, and in this case it will be more accurate to use the second expression with this direct estimate than the first expression.

We conditioned the probability distribution on a given number of extant species at a fixed time point in the past, and in practice we used this conditioning to force the expected number of species to be 1 at the stem age, or 2 at the crown age of the group. Stem age estimates are not always available and may be less accurate than crown ages. In this case, the crown condition should be used. However when stem ages are available and reliable, using the stem condition should be preferred, as forcing the existence of exactly two lineages at the crown age ignores extinctions that might have happened between the stem and the crown age. In addition, the stem age is insensitive to extinctions and undersampling while the crown age can be underestimated when there are missing species.

We have shown that the DTT curves obtained with the probabilistic approach are more accurate than those obtained with LTT plots and the deterministic approach of Morlon et al. (2011); in particular, they are less biased towards an underestimation of past species richness. They also avoid some misbehaviors of the deterministic approach in some specific cases, as illustrated here with the Didelphidae. In addition, they are more robust to undersampling, and not deeply affected by reasonable departures from models’ assumptions. Finally, they offer the notable advantage of providing confidence intervals around diversity estimates. The confidence intervals computed here do not account for the uncertainty in rate estimates. In future developments, one could imagine incorporating such uncertainty by replacing the probability expressions in the computation of the DTT with their averages over posterior rate estimates. This would provide confidence intervals over the data, as opposed to confidence intervals given the parameters, as computed here.

While the approach presented here improves on previous approaches, it has some limitations. First, the accuracy of the diversity estimates critically depends on the accuracy of the speciation and extinction rate estimates, which is not always high; for example, extinction rates estimated in phylogenetic studies are often unrealistically low, and more generally the diversification model selected is not always accurate. Our measure of error between simulated and estimated diversity reflects in large part this error in rate estimates. At the same time, the improved performance of this approach as compared to the deterministic one shows that the approach better handles such rate uncertainties, probably thanks to the conditioning of the number of species at the origin of the group.

There are many possible sources of biases not investigated here and room for improvement. For example, we used a likelihood formula that assumes uniform sampling; in reality, the sampling is most likely not uniform, which might lead to biased rate estimates (Höhna et al., 2011). In principle, one could use likelihoods accounting for other sampling schemes (Höhna et al., 2011) to estimate rates through time, and Equation 1 to deduce DTT plots. Similarly, our equations can in principle be used in combination with any other diversification model that provides estimates of diversification rates through time, such as piecewise constant rate estimates with mass extinction events (Stadler, 2011), or models accounting for environmental dependencies (Condamine et al., 2013; Lewitus and Morlon, 2017). We did not implement this here and we only investigated a subset of the potential biases. In general, we expect the approach to perform the best in situations when the rates are well estimated.

Under most model scenarios, we found that archaeobatrachian diversity peaked in the deep past, about 160 million years ago, and then gradually declined until more recently. Yet remnants of this early frog diversity continue to the present day in the small families of Archaeobatrachia, which – according to our results – add up to a total number of species that is much smaller than it used to be. Additionally, a young, high-diversity group (Neobatrachia) is nested within this older, low-diversity group. This pattern is relatively common and found in such groups as Lepidosauria (tuataras vs. squamates), Serpentes (Colubroidea vs. other snake clades), and whales. A common hypothesis for the decline in diversity of the old, species-poor groups is that it relates to the rise of the younger groups, but it is unclear whether that happened in frogs. According to our results, Archaeobatrachia shows a gradual loss of species starting from about 160 Ma, which is long before the younger families started to rise in diversity, particularly near the Cretaceous-Tertiary mass extinction 66 Ma (Feng et al., 2017). A recent phylogenomic analysis of major frog lineages (Feng et al., 2017) suggests younger divergence times than the ones from Pyron (2014) that we used here. How these new divergence estimates will affect our findings is unclear. However, because divergence times in Feng et al. (2017) were estimated to be more recent mainly in young families, we can speculate that the lag between the decline of Archaeobatrachia and the rise of younger families is underestimated, rather than overestimated, in our study.

Determining what caused the decline of Archaeobatrachia is difficult without more detailed fossil data. However, the pattern of a symmetric rise and fall in diversity, especially coupled with the rise in diversity of a nested clade, suggests competitive replacement (Sepkoski Jr et al., 2000; Silvestro et al., 2015; Vermeij, 1987). Competitive replacement is often presented as part of the Red Queen hypothesis of Van Valen (1973) (see also Liow et al. 2011), which states that an organism’s environment – particularly its biotic environment – is always changing, and if members of a clade do not keep up with the constant change, the clade will go extinct. This implies that species diversity in clades will rise and fall over time, particularly if a similar group of organisms (e.g. members of that clade) does adapt and outcompetes the species that do not change. Whether this happened for Archaeobatrachia is unclear. Biogeographic analyses show that Neobatrachia most likely originated during the splitting of Pangaea in two, with most archaeobatrachian lineages staying on Laurasia, and Neobatrachia originating on Gondwana (Feng et al., 2017; Roelants and Bossuyt, 2005). Subsequent neobatrachian colonization of North America and Eurasia likely happened much later than our inferred declined of Archaeobatrachia (Fig. 6; Feng et al. (2017)). Moreover, our analyses show that the decline of Archaeobatrachia is associated with a failure to speciate rather than with increased extinction rates, a pattern that has previously been observed in mammals but whose causes are not well understood (Quental and Marshall, 2013). Such analyses of diversity trajectories have been performed mostly using fossil data (Quental and Marshall, 2013; Sepkoski Jr et al., 2000; Silvestro et al., 2015). The method we develop and apply here will allow investigators to address such questions in other groups, particularly those without an extensive fossil record.

Following the mathematical approach adopted here, one could condition the probability distribution for the number of extant species over time on a given number of species at more than a single fixed time point in the past, as we do here at the root. This could be useful if we had a good estimate, for example from the fossil record, of the number of species at specific times in the past (e.g. periods when preservation was particularly good). This could provide a well-needed approach for integrating phylogenetic and fossil information in order to improve our understanding of past diversity dynamics (Condamine et al., 2013; Heath et al., 2014; Morlon, 2014).

We expect that the approach outlined here will be useful for more than just estimating the diversity-through-time curve of particular clades. For example, there is increasing interest in understanding the role of clade-clade competition in diversification (Silvestro et al., 2015), but this question has not been addressed in groups with a poor fossil record, due to a lack of appropriate phylogenetic comparative approaches. One could test if and how diversification in one clade (clade A) is influenced by the number of species in a putatively competing clade (clade B) by first estimating the DTT curve of clade B using the approach developed here, and next evaluating if diversification in clade A has been influenced by species richness in clade B using environment-dependent models of diversification (Condamine et al., 2013; Morlon, 2014), with species richness in clade B used as the “environment”. These models are already implemented in RPANDA (Morlon et al., 2016).

Our approach could also be used to improve so-called diversity-dependent models of phenotypic evolution, in which the rate of phenotypic evolution depends on the number of extant species in a clade (Mahler et al., 2010; Weir and Mursleen, 2013). These models have been developed in the context of adaptive radiations (Simpson, 1955), with the underlying idea that evolution should slow down as ecological niches are filled during adaptive radiations (Moen and Morlon, 2014; Schluter, 2000). Hence, diversity-dependent models of phenotypic evolution have been used as a test of adaptive radiations (Mahler et al., 2010; Weir and Mursleen, 2013). In the absence of a better option, these models have used the number of reconstructed lineages (that is, LTT plots) as a proxy for the number of extant species at a given time in the past, thus ignoring extinction (Mahler et al., 2010; Weir and Mursleen, 2013). This has been shown to lead to an underestimation of diversity-dependent effects (Drury et al., 2016). As an alternative to using reconstructed lineages, one could use our diversity-through-time estimates, which we have shown are more accurate. This should significantly improve the performance of these models. Once accurate DTT curves have been computed, one can analyze if and how species richness influences the rate at which phenotypes evolve using environment-dependent models of phenotypic evolution (Clavel and Morlon, 2017), with species richness used as the “environment”. These models are also already implemented in RPANDA (Morlon et al., 2016). A similar approach could also be used to test clade-clade co-evolutionary scenarios, such as the rate of phenotypic evolution in a clade (e.g. evolution of chemical defences in plants) being influenced by the number of species in the interacting clade (e.g. the herbivores that feed on plants).

## SUPPLEMENTARY MATERIAL

Data available from the Dryad Digital Repository:

## ACKNOWLEDGMENTS

We thank members of the Morlon lab for comments on the manuscript.

## References

Alfaro, M., F. Santini, C. Brock, H. Alamillo, A. Dornburg, D. L Rabosky, G. Carnevale, and L. Harmon. 2009. Nine exceptional radiations plus high turnover explain species diversity in jawed vertebrates. Proc. Natl. Acad. Sci. 106:13410–13414.

AmphibiaWeb. 2016. Amphibiaweb: information on amphibian biology and conservation. http://amphibiaweb.org [Online; accessed 1 August 2016].

Apesteguía, S. and F. E. Novas. 2003. Large Cretaceous sphenodontian from Patagonia provides insight into lepidosaur evolution in Gondwana. Nature 425:609–612.

Beaulieu, J. M. and B. C. O’Meara. 2015. Extinction can be estimated from moderately sized molecular phylogenies. Evolution 69:1036–1043.

Clavel, J. and H. Morlon. 2017. Accelerated body size evolution during cold climatic periods in the Cenozoic. Proc. Natl. Acad. Sci. 114:4183–4188.

Condamine, F. L., J. Rolland, and H. Morlon. 2013. Macroevolutionary perspectives to environmental change. Ecol. Lett. 16:72–85.

Drury, J., J. Clavel, M. Manceau, and H. Morlon. 2016. Estimating the effect of competition on trait evolution using maximum likelihood inference. Syst. Biol. 65:700–710.

Duellman, W. E. 1975. On the classification of frogs. Occasional Papers of the Museum of Natural History, The University of Kansas 42:1–15.

Etienne, R. S., B. Haegeman, T. Stadler, T. Aze, P. N. Pearson, A. Purvis, and A. B. Phillimore. 2011. Diversity-dependence brings molecular phylogenies closer to agreement with the fossil record. Proc. R. Soc. B 279:1300–1309.

Ezard, T. H., T. Aze, P. N. Pearson, and A. Purvis. 2011. Interplay between changing climate and species’ ecology drives macroevolutionary dynamics. Science 332:349–351.

Ezard, T. H. and A. Purvis. 2016. Environmental changes define ecological limits to species richness and reveal the mode of macroevolutionary competition. Ecol. Lett. 19:899–906.

Faurby, S. and J.-C. Svenning. 2015. A species-level phylogeny of all extant and late Quaternary extinct mammals using a novel heuristic-hierarchical Bayesian approach. Mol. Phyl. Evol. 84:14–26.

Feng, Y.-J., D. C. Blackburn, D. Liang, D. M. Hillis, D. B. Wake, D. C. Cannatella, and P. Zhang. 2017. Phylogenomics reveals rapid, simultaneous diversification of three major clades of Gondwanan frogs at the Cretaceous-Paleogene boundary. Proc. Natl. Acad. of Sci. 114:E5864–E5870.

Foote, M., J. S. Crampton, A. G. Beu, B. A. Marshall, R. A. Cooper, P. A. Maxwell, and I. Matcham. 2007. Rise and fall of species occupancy in Cenozoic fossil mollusks. Science 318:1131–1134.

Ford, L. S. and D. C. Cannatella. 1993. The major clades of frogs. Herpetol. Monogr. Pages 94–117.

Harmon, L. J. 2012. An inordinate fondness for eukaryotic diversity. PLoS Biol. 10:e1001382.

Heath, T. A., J. P. Huelsenbeck, and T. Stadler. 2014. The fossilized birth-death process for coherent calibration of divergence-time estimates. Proc. Natl. Acad. Sci. 111:E2957–E2966.

Höhna, S., T. Stadler, F. Ronquist, and T. Britton. 2011. Inferring speciation and extinction rates under different sampling schemes. Molecular Biology and Evolution 28:2577–2589.

Jetz, W. and R. A. Pyron. 2018. The interplay of past diversification and evolutionary isolation with present imperilment across the amphibian tree of life. Nature Ecology & Evolution 2:850–858.

Jones, M. E., A. J. Tennyson, J. P. Worthy, S. E. Evans, and T. H. Worthy. 2009. A sphenodontine (Rhynchocephalia) from the Miocene of New Zealand and palaeobiogeography of the tuatara (*Sphenodon*). Proc. R. Soc. B 276:1385–1390.

Kendall, D. G. 1948. On the generalized “birth-and-death” process. Ann. Math. Stat. Pages 1–15.

Lambert, A. 2018. The coalescent of a sample from a binary branching process. Theoret. Popul. Biol. 122:30–35.

Lewitus, E., L. Bittner, S. Malviya, C. Bowler, and H. Morlon. 2018. Clade-specific diversification dynamics of marine diatoms since the jurassic. Nature Ecology & Evolution 2:1715–1723.

Lewitus, E. and H. Morlon. 2017. Detecting environment-dependent diversification from phylogenies: A simulation study and some empirical illustrations. Syst. Biol. 67:576–593.

Liow, L. H., T. Reitan, and P. G. Harnik. 2015. Ecological interactions on macroevolutionary time scales: clams and brachiopods are more than ships that pass in the night. Ecol. Lett. 18:1030–1039.

Liow, L. H., L. Van Valen, and N. C. Stenseth. 2011. Red Queen: from populations to taxa and communities. Trends Ecol. Evol. 26:349–358.

Louca, S., P. M. Shih, M. W. Pennell, W. W. Fischer, L. W. Parfrey, and M. Doebeli. 2018. Bacterial diversification through geological time. Nature Ecology & Evolution 2:1458.

Mahler, D. L., L. J. Revell, R. E. Glor, and J. B. Losos. 2010. Ecological opportunity and the rate of morphological evolution in the diversification of Greater Antillean anoles. Evolution 64:2731–2745.

Marshall, C. R. and T. B. Quental. 2016. The uncertain role of diversity dependence in species diversification and the need to incorporate time-varying carrying capacities. Phil. Ttans. R. Soc. B 371:20150217.

Moen, D. and H. Morlon. 2014. Why does diversification slow down? Trends Ecol. Evol. 29:190–197.

Moore, B. R., S. Hohna, M. R. May, B. Rannala, and J. P. Huelsenbeck. 2016. Critically evaluating the theory and performance of bayesian analysis of macroevolutionary mixtures. Proceedings of the National Academy of Sciences 113:9569–9574.

Morlon, H. 2014. Phylogenetic approaches for studying diversification. Ecol. Lett. 17:508–525.

Morlon, H., B. D. Kemps, J. B. Plotkin, and D. Brisson. 2012. Explosive radiation of a bacterial species group. Evolution 66:2577–2586.

Morlon, H., E. Lewitus, F. L. Condamine, M. Manceau, J. Clavel, and J. Drury. 2016. RPANDA: an R package for macroevolutionary analyses on phylogenetic trees. Meth. Ecol. Evol. 7:589–597.

Morlon, H., T. L. Parsons, and J. B. Plotkin. 2011. Reconciling molecular phylogenies with the fossil record. Proc. Natl. Acad. Sci. 108:16327–16332.

Nee, S., A. O. Mooers, and P. H. Harvey. 1992. Tempo and mode of evolution revealed from molecular phylogenies. Proc. Natl. Acad. Sci. 89:8322–8326.

Paradis, E. 2011. Time-dependent speciation and extinction from phylogenies: A least squares approach. Evolution 65:661–672.

Pennell, M. W. and L. J. Harmon. 2013. An integrative view of phylogenetic comparative methods: connections to population genetics, community ecology, and paleobiology. Ann. N. Y. Acad. Sci. 1289:90–105.

Pough, F., R. Andrews, M. Crump, A. Savitzsky, K. Wells, and M. Brandley. 2015. Herpetology. Sinauer Associates.

Pyron, R. A. 2014. Biogeographic analysis reveals ancient continental vicariance and recent oceanic dispersal in amphibians. Syst. Biol. 63:779–797.

Pyron, R. A. and J. J. Wiens. 2011. A large-scale phylogeny of Amphibia including over 2800 species, and a revised classification of extant frogs, salamanders, and caecilians. Mol. Phyl. Evol. 61:543–583.

Quental, T. B. and C. R. Marshall. 2010. Diversity dynamics: molecular phylogenies need the fossil record. Trends Ecol. Evol. 25:434–441.

Quental, T. B. and C. R. Marshall. 2013. How the Red Queen drives terrestrial mammals to extinction. Science 341:290–292.

Quince, C., T. P. Curtis, and W. T. Sloan. 2008. The rational exploration of microbial diversity. The ISME Journal 2:997–1006.

Rabosky, D. L. 2010. Extinction rates should not be estimated from molecular phylogenies. Evolution 64:1816–1824.

Rabosky, D. L. 2014. Automatic detection of key innovations, rate shifts, and diversity-dependence on phylogenetic trees. PloS One 9:e89543.

Rabosky, D. L. and U. Sorhannus. 2009. Diversity dynamics of marine planktonic diatoms across the Cenozoic. Nature 457:183–186.

Ricklefs, R. E. 2007. Estimating diversification rates from phylogenetic information. Trends Ecol. Evol. 22:601–610.

Roelants, K. and F. Bossuyt. 2005. Archaeobatrachian paraphyly and Pangaean diversification of crown-group frogs. Syst. Biol. 54:111–126.

Sanderson, M. J. 1996. How many taxa must be sampled to identify the root node of a large clade? Syst. Biol. 45:168–173.

Schluter, D. 2000. The ecology of adaptive radiation. OUP Oxford.

Sepkoski Jr, J. J., F. K. McKinney, and S. Lidgard. 2000. Competitive displacement among post-Paleozoic cyclostome and cheilostome bryozoans. Paleobiology 26:7–18.

Silvestro, D., A. Antonelli, N. Salamin, and T. B. Quental. 2015. The role of clade competition in the diversification of North American canids. Proc. Natl. Acad. Sci. 112:8684–8689.

Simpson, G. G. 1955. Major features of evolution. Columbia University Press: New York.

Stadler, T. 2011. Mammalian phylogeny reveals recent diversification rate shifts. Proc. Natl. Acad. Sci. 108:6187–6192.

Stadler, T. 2013. Recovering speciation and extinction dynamics based on phylogenies. J. Evol. Biol. 26:1203–1219.

Stadler, T. 2015. TreeSim: Simulating Phylogenetic Trees. R package version 2.2.

Steeman, M. E., M. B. Hebsgaard, R. E. Fordyce, S. Y. Ho, D. L. Rabosky, R. Nielsen, C. Rahbek, H. Glenner, M. V. Sørensen, and E. Willerslev. 2009. Radiation of extant cetaceans driven by restructuring of the oceans. Syst. Biol. 58:573–585.

Uetz, P., J. Hošek, and J. Hallermann. 2018. The reptile database. http://www.reptile-database.org [Online; accessed 26 October 2018].

Van Valen, L. 1973. A new evolutionary law. Evol. Theor.

Vermeij, G. J. 1987. Evolution and escalation: an ecological history of life. Princeton University Press.

Weir, J. T. and S. Mursleen. 2013. Diversity-dependent cladogenesis and trait evolution in the adaptive radiation of the auks (Aves: Alcidae). Evolution 67:403–416.

