## Appendices for "Estimating Diversity Through Time using Molecular Phylogenies: Old and Species-Poor Frog Families are the Remnants of a Diverse Past"

Billaud, O., Moen, D. S., Parsons, T. L., Morlon, H.

November 29, 2018

In this appendix, we present the derivations of the results presented in the Methods section. We are considering the number of species in a conditioned time-inhomogeneous birth-death model of cladogenesis, which is equivalent to considering the number of individuals in a conditioned time-inhomogeneous birth and death process. We thus begin by recalling the relevant results for the latter. Given these, we apply Bayes' theorem to obtain the one-dimensional distributions of the process conditioned on the number of species in the present and at some fixed point in the past where we know the number of species present (*e.g.* stem age or crown age). Assuming that each extant species is sampled with probability $f$, we can use those distributions to obtain the one-dimensional distributions for the process conditioned on the number of observed species in terms of derivatives of the generating function for the process conditioned on the total number of species at the present. We then show how those derivatives may be obtained analytically. Finally, we present several additional results for these conditioned processes that were not directly used in the main text, namely an approach for computing confidence intervals, an analytical expression for the expected number of lines through time, and rates for a Markov chain that may be simulated to give sample diversity through time plots with the appropriate stem or crown age and number of (observed) species in the present.

#### A Preliminaries for the Time-Inhomogeneous Birth and Death Process

We will assume $N(t)$ is a time-inhomogeneous birth-and-death process, with time varying transition intensities

$$q_{n,n+1}(t) = \lambda(t) \text{ and } q_{n,n-1}(t) = \mu(t).$$

so the generator of the process $N(t)$ is

$$\begin{aligned}(\mathcal{G}_t f)(n) &= \lim_{h\downarrow 0} \frac{\mathbb{E}\left[f(N(t+h))|N(t)=n\right] - f(n)}{h} \\ &= \lambda(t)n\left(f(n+1)-f(n)\right) + \mu(t)n\left(f(n-1)-f(n)\right)\end{aligned}$$

for functions $f : \mathbb{N}_0 \to \mathbb{R}$.

Let $F_x(z,s,t)$ be the probability generating function for $N(t)$, so

$$F_x(z,s,t) := \mathbb{E}\left[ z^{N(t)} \middle| N(s) = x \right]. \tag{A.1}$$

Then, $F_x = F^x$, where $F(z,s,t) := F_1(z,s,t)$ is the unique solution to the backward equation

$$\frac{\partial F}{\partial t} = -\mathcal{G}_t F.$$

**Theorem 1** (Kendall (1948))**.** *Let*

$$q(s,t) := \frac{\int_s^t e^{-\int_s^\tau \lambda(u)-\mu(u)\,du}\mu(\tau)\,d\tau}{1+\int_s^t e^{-\int_s^\tau \lambda(u)-\mu(u)\,du}\mu(\tau)\,d\tau},$$

*and*

$$\eta(s,t) := \frac{\int_s^t e^{\int_\tau^t \lambda(u)-\mu(u)\,du}\lambda(\tau)\,d\tau}{1+\int_s^t e^{\int_\tau^t \lambda(u)-\mu(u)\,du}\lambda(\tau)\,d\tau},$$

*Then,*

$$F(z,s,t) = \frac{q(s,t)+(1-q(s,t)-\eta(s,t))z}{1-\eta(s,t)z}, \tag{A.2}$$

*and, if* $P_n(s,t) := \mathbb{P}\{N(t)=n|N(s)=1\}$, *we have*

$$F(z,s,t) = \sum_{n=0}^{\infty} P_n(s,t) z^n$$

*for*

$$P_n(s,t) = \begin{cases} q(s,t) & \textit{if } n=0 \\ (1-q(s,t))(1-\eta(s,t))\eta(s,t)^{n-1} & \textit{if } n \geq 1. \end{cases}$$

We also note, for further use, that

$$\begin{aligned}\mathbb{P}(N(t)=n|N(s)=x) &= [z^n](F(z,s,t))^x \\ &= \sum_{n_1+\cdots+n_x=n} P_{n_1}(s,t)\cdots P_{n_x}(s,t).\end{aligned}$$

Now, if $k$ of the variables $n_j$ are 0, so that the remaining $m-k$ sum to $n$, then we have

$$\begin{aligned}&P_{n_1}(s,t)\cdots P_{n_x}(s,t) \\ &\qquad = q(s,t)^k(1-q(s,t))^{x-k}(1-\eta(s,t))^{x-k}\eta(s,t)^{n-x+k}.\end{aligned}$$

Further, there are $\binom{x}{k}$ ways of picking $k$ values $n_j$ to be 0, and $\binom{n-1}{x-k-1}$ ways that the remaining $x-k$ values $n_j$ can sum to $n$, so

$$\begin{aligned}\mathbb{P}(N(t)=n|N(s)=x) &= \sum_{k=0}^{x-1}\binom{x}{k}\binom{n-1}{x-k-1} q(s,t)^k(1-q(s,t))^{x-k}(1-\eta(s,t))^{x-k}\eta(s,t)^{n-x+k} \\ &= (1-q(s,t))^x(1-\eta(s,t))^x\eta(s,t)^{n-x}\sum_{k=0}^{x-1}\binom{x}{k}\binom{n-1}{x-k-1}\left(\frac{q(s,t)\eta(s,t)}{(1-q(s,t))(1-\eta(s,t))}\right)^k.\end{aligned}$$

### B Distribution for the Conditioned Process

We are interested in the branching process conditioned on initial and final conditions. We express these going forward in time, so that $t = 0$ is the time of the most recent common ancestor, and our sample will be taken at time $T_{\mathrm{mrca}}$, which is the present. We assume that $N(s) = x$ for some $0 \leq s \leq T_{\mathrm{mrca}}$ and that $N(T_{\mathrm{mrca}}) = n$.

We compute the probability conditional on $N(s) = x$ and $N(T_{\mathrm{mrca}}) = n$. This is done via Bayes' Theorem, which tells us that, given events $A$ and $B$,

$$\mathbb{P}(B|A) = \frac{\mathbb{P}(B \cap A)}{\mathbb{P}(A)} = \frac{\mathbb{P}(A|B)\,\mathbb{P}(B)}{\mathbb{P}(A)}$$

In particular, if the space of all possible events, $\Omega$ can be written as countable union of disjoint sets, *i.e.*, $\Omega = \bigcup_j B_j$, $B_i \cap B_j = \emptyset$ for $i \neq j$, then

$$\mathbb{P}(B|A) = \frac{\mathbb{P}(A|B)\,\mathbb{P}(B)}{\sum_j \mathbb{P}(A|B_j)\,\mathbb{P}(B_j)}.$$

We shall use both forms.

So to start, we observe that for $s \leq t \leq T_{\mathrm{mrca}}$,

$$\begin{aligned}
&\mathbb{P}(N(t) = m | N(s) = x, N(T_{\mathrm{mrca}}) = n) \\
&\qquad = \frac{\mathbb{P}(N(t) = m, N(T_{\mathrm{mrca}}) = n | N(s) = x)}{\mathbb{P}(N(T_{\mathrm{mrca}}) = n | N(s) = x)} \\
&\qquad = \frac{\mathbb{P}(N(T_{\mathrm{mrca}}) = n | N(t) = m, N(s) = x)\,\mathbb{P}(N(t) = m | N(s) = x)}{\mathbb{P}(N(T_{\mathrm{mrca}}) = n | N(s) = x)}.
\end{aligned}$$

Now, by the Markov property,

$$\mathbb{P}(N(T_{\mathrm{mrca}}) = n | N(t) = m, N(s) = x) = \mathbb{P}(N(T_{\mathrm{mrca}}) = n | N(t) = m),$$

so the expression above simplifies to

$$\begin{aligned}
&\mathbb{P}(N(t) = m | N(s) = x, N(T_{\mathrm{mrca}}) = n) \\
&\qquad = \frac{\mathbb{P}(N(T_{\mathrm{mrca}}) = n | N(t) = m)\,\mathbb{P}(N(t) = m | N(s) = x)}{\mathbb{P}(N(T_{\mathrm{mrca}}) = n | N(s) = x)},
\end{aligned}$$

which can be explicitly computed using Kendall's results above.

### C Conditioning on the Number of Observed Lines

We now compute the probability conditional on having observed $N_{\mathrm{obs}}(T_{\mathrm{mrca}}) = l$ lineages at the present, assuming that we have a probability $f$ of observing a line, given that it is alive.

The number of observed lines, conditional on $N(T_{\mathrm{mrca}})$ is binomially distributed:

$$\mathbb{P}(N_{\mathrm{obs}}(T_{\mathrm{mrca}}) = l | N(T_{\mathrm{mrca}}) = n) = \binom{n}{l} f^l (1-f)^{n-l}.$$

Again, we apply Bayes' Theorem to conclude

$$\begin{aligned}
\mathbb{P}\left(N(T_{\text{mrca}})=n|N_{\text{obs}}(T_{\text{mrca}})=l,N(s)=x\right) &= \frac{\mathbb{P}\left(N(T_{\text{mrca}})=n,N_{\text{obs}}(T_{\text{mrca}})=l|N(s)=x\right)}{\mathbb{P}\left(N_{\text{obs}}(T_{\text{mrca}})=l|N(s)=x\right)} \\
&= \frac{\mathbb{P}\left(N_{\text{obs}}(T_{\text{mrca}})=l|N(T_{\text{mrca}})=n,N(s)=x\right)\mathbb{P}\left(N(T_{\text{mrca}})=n|N(s)=x\right)}{\sum_{j=1}^{\infty}\mathbb{P}\left(N_{\text{obs}}(T_{\text{mrca}})=l|N(T_{\text{mrca}})=j,N(s)=x\right)\mathbb{P}\left(N(T_{\text{mrca}})=j|N(s)=x\right)} \\
&= \frac{\mathbb{P}\left(N_{\text{obs}}(T_{\text{mrca}})=l|N(T_{\text{mrca}})=n\right)\mathbb{P}\left(N(T_{\text{mrca}})=n|N(s)=x\right)}{\sum_{j=1}^{\infty}\mathbb{P}\left(N_{\text{obs}}(T_{\text{mrca}})=l|N(T_{\text{mrca}})=j\right)\mathbb{P}\left(N(T_{\text{mrca}})=j|N(s)=x\right)} \\
&= \frac{\binom{n}{l}f^l(1-f)^{n-l}\mathbb{P}\left(N(T_{\text{mrca}})=n|N(s)=x\right)}{\sum_{j=l}^{\infty}\binom{j}{l}f^l(1-f)^{j-l}\mathbb{P}\left(N(T_{\text{mrca}})=j|N(s)=x\right)} \\
&= \frac{(n)_l(1-f)^n\mathbb{P}\left(N(T_{\text{mrca}})=n|N(s)=x\right)}{\sum_{j=l}^{\infty}(j)_l(1-f)^j\mathbb{P}\left(N(T_{\text{mrca}})=j|N(s)=x\right)},
\end{aligned}$$

where $(n)_l = \frac{n!}{(n-l)!}$ is the falling factorial.

Now,

$$\begin{aligned}
z^l\frac{\partial^l}{\partial z^l}F_x(z,s,T_{\text{mrca}}) &= z^l\frac{\partial^l}{\partial z^l}\sum_{j=0}^{\infty}\mathbb{P}\left(N(T_{\text{mrca}})=j|N(s)=x\right)z^j \\
&= z^l\sum_{j=l}^{\infty}(j)_l\mathbb{P}\left(N(T_{\text{mrca}})=j|N(s)=x\right)z^{j-l} \\
&= \sum_{j=l}^{\infty}(j)_l\mathbb{P}\left(N(T_{\text{mrca}})=j|N(s)=x\right)z^j,
\end{aligned}$$

so the infinite sum in the denominator is

$$(1-f)^l\frac{\partial^l F_x}{\partial z^l}(1-f,s,T_{\text{mrca}}).$$

Recalling that $F_x = F^x$, we can again use Kendall's results. We note that Kendall's expression for the generating series, (A.2), has a unique pole at $z=\frac{1}{\eta(s,t)}$, and thus has radius of convergence $R=\frac{1}{\eta(s,t)}\geq 1$, and may be evaluated at $z=1-f$.

Now,

$$\begin{aligned}
&\mathbb{P}\left(N(t)=m|N(s)=x,N_{\text{obs}}(T_{\text{mrca}})=l\right) \\
&\quad= \sum_{n=l}^{\infty}\mathbb{P}\left(N(t)=m,N(T_{\text{mrca}})=n|N(s)=x,N_{\text{obs}}(T_{\text{mrca}})=l\right) \\
&\quad= \sum_{n=l}^{\infty}\mathbb{P}\left(N(t)=m|N(s)=x,N_{\text{obs}}(T_{\text{mrca}})=l,N(T_{\text{mrca}})=n\right) \\
&\qquad\qquad\times\mathbb{P}\left(N(T_{\text{mrca}})=n|N_{\text{obs}}(T_{\text{mrca}})=l,N(s)=x\right).
\end{aligned}$$

Conditioning on the number observed gives no additional information from conditioning on the number actually present, so the latter is

$$\sum_{n=l}^{\infty} \mathbb{P}\left(N(t)=m | N(s)=x, N(T_{\mathrm{mrca}})=n\right) \\ \times \mathbb{P}\left(N(T_{\mathrm{mrca}})=n | N_{\mathrm{obs}}(T_{\mathrm{mrca}})=l, N(s)=x\right).$$

Combining our previous results, we then have

$$\mathbb{P}\left(N(t)=m | N(s)=x, N_{\mathrm{obs}}(T_{\mathrm{mrca}})=l\right) \\ = \frac{\sum_{n=l}^{\infty}(n)_l(1-f)^n \mathbb{P}\left(N(T_{\mathrm{mrca}})=n | N(t)=m\right) \mathbb{P}\left(N(t)=m | N(s)=x\right)}{(1-f)^l \frac{\partial^l F_x}{\partial z^l}(1-f, s, T_{\mathrm{mrca}})}.$$

Finally, we observe as before that we can collapse the infinite sum as before to get

$$\sum_{n=l}^{\infty}(n)_l(1-f)^n \mathbb{P}\left(N(T_{\mathrm{mrca}})=n | N(t)=m\right) = (1-f)^l \frac{\partial^l F_m}{\partial z^l}(1-f, t, T_{\mathrm{mrca}})$$

so that

$$\mathbb{P}\left(N(t)=m | N(s)=x, N_{\mathrm{obs}}(T_{\mathrm{mrca}})=l\right) = \frac{\frac{\partial^l F_m}{\partial z^l}(1-f, t, T_{\mathrm{mrca}})}{\frac{\partial^l F_x}{\partial z^l}(1-f, s, T_{\mathrm{mrca}})} \mathbb{P}\left(N(t)=m | N(s)=x\right).$$

Taking $l=n$ and $f=1$, we see that all but the lowest order term vanishes and

$$\mathbb{P}\left(N(T_{\mathrm{mrca}})=n | N(t)=m\right) = \frac{1}{n!} \frac{\partial^n F_m}{\partial z^n}(0, t, T_{\mathrm{mrca}}) \tag{A.3}$$

and

$$\mathbb{P}\left(N(T_{\mathrm{mrca}})=n | N(s)=x\right) = \frac{1}{n!} \frac{\partial^n F_x}{\partial z^n}(0, s, T_{\mathrm{mrca}}) \tag{A.4}$$

so the expression for the probability conditioned on the total number of extant species becomes

$$\mathbb{P}\left(N(t)=m | N(s)=x, N(T_{\mathrm{mrca}})=n\right) = \frac{\frac{\partial^n F_m}{\partial z^n}(0, t, T_{\mathrm{mrca}})}{\frac{\partial^n F_x}{\partial z^n}(0, s, T_{\mathrm{mrca}})} \mathbb{P}\left(N(t)=m | N(s)=x\right),$$

which is consistent with our previous expression.

Finally, we can use the above to write the probability distribution of the process conditioned on the number of observed lines completely in terms of the generating function:

$$\mathbb{P}\left(N(t)=m | N(s)=x, N_{\mathrm{obs}}(T_{\mathrm{mrca}})=l\right) = \frac{1}{m!} \frac{\frac{\partial^l F_m}{\partial z^l}(1-f, t, T_{\mathrm{mrca}})}{\frac{\partial^l F_x}{\partial z^l}(1-f, s, T_{\mathrm{mrca}})} \frac{\partial^m F_x}{\partial z^m}(0, s, t).$$

### D Evaluating the Derivatives

To evaluate the derivatives $\frac{\partial^l F_m}{\partial z^l}(1-f, t, T_{\mathrm{mrca}})$, we exploit the fact that if $f$ is analytic at $z = a$ and $R$ is the radius of convergence of $f$ at $a$, then for $|z-a| < R$, we have

$$f(z) = \sum_{n=0}^{\infty} \frac{f^{(n)}(a)}{n!}(z-a)^n$$

(here, $f^{(n)}(z)$ is the $n^{\mathrm{th}}$ derivative of $f$), and, moreover, this expansion is unique, *i.e.*, if we can write

$$f(z) = \sum_{n=0}^{\infty} c_n (z-a)^n,$$

then $c_n = \frac{f^{(n)}(a)}{n!}$. We will do so by finding a series expansion for $F^m(z,t,s)$ about $a := 1-f$, using Kendall's closed form expression for $F(z,s,t)$, which may be done using the binomial theorem and the binomial series. For the latter, we recall that the binomial coefficients can be defined for all $\alpha \in \mathbb{C}$ by

$$\binom{\alpha}{k} := \frac{(\alpha)_k}{k!},$$

and then, for $|z| < 1$,

$$(1+z)^\alpha = \sum_{k=0}^{\infty} \binom{\alpha}{k} z^k.$$

To use this, we observe that

$$\begin{aligned}
F^m(z,s,t) &= \left(\frac{q(s,t) + (1 - q(s,t) - \eta(s,t))z}{1 - \eta(s,t)z}\right)^m \\
&= ((q(s,t) + a(1 - q(s,t) - \eta(s,t))) + (1 - q(s,t) - \eta(s,t))(z-a))^m \\
&\quad \times ((1 - a\eta(s,t)) - \eta(s,t)(z-a))^m \\
&= (A(s,t) + B(s,t)\zeta)^m \, (C(s,t) - \eta(s,t)\zeta)^{-m}
\end{aligned}$$

where, for simplicity, we set $A(s,t) := q(s,t) + a(1 - q(s,t) - \eta(s,t))$, $B(s,t) := 1 - q(s,t) - \eta(s,t)$, $C(s,t) := 1 - a\eta(s,t)$, and $\zeta := z - a$.

If we expand the latter in a series in $\zeta$, then the coefficient of $\zeta^l$ will be $\frac{1}{l!}\frac{\partial^l F_m}{\partial z^l}(1-f,s,t)$. To do so, we apply the binomial theorem to the first term in the product, and expand the second as a binomial series:

$$\begin{aligned}
&(A(s,t) + B(s,t)\zeta)^m \, (C(s,t) - \eta(s,t)\zeta)^{-m} \\
&\qquad = \frac{1}{C^m(s,t)} (A(s,t) + B(s,t)\zeta)^m \left(1 - \frac{\eta(s,t)}{C(s,t)}\zeta\right)^{-m}
\end{aligned}$$

For the latter, we note that the binomial series converges if $\left|\frac{\eta(s,t)}{C(s,t)}\zeta\right| < 1$, *i.e.*, if

$$|z-a| < \frac{C(s,t)}{\eta(s,t)},$$

which can be satisfied for fixed $s,t$ for $z$ sufficiently close to $a = 1-f$:

$$\frac{C(s,t)}{\eta(s,t)} = \frac{1}{\eta(s,t)} - a \geq 1 - a = f.$$

Because we are only concerned with the coefficients, we can choose $z$ arbitrarily close to $a$ as needed. Then,

$$(A(s,t) + B(s,t)\zeta)^m = \sum_{j=0}^{m} \binom{m}{j} A(s,t)^{m-j} B(s,t)^j \zeta^j,$$

and

$$\frac{1}{C^m(s,t)} \left(1 - \frac{\eta(s,t)}{C(s,t)}\zeta\right)^{-m} = \frac{1}{C^m(s,t)} \sum_{k=0}^{\infty} \binom{-m}{k} (-1)^k \frac{\eta^k(s,t)}{C^k(s,t)} \zeta^k.$$

We seek the coefficient of $\zeta^l$ in the product, which is the sum over all terms with $j+k=l$, *i.e.*,

$$\begin{aligned}
&\frac{1}{C^m(s,t)} \sum_{\substack{j+k=l \\ j\leq m}} (-1)^k \binom{m}{j} \binom{-m}{k} \frac{A^{m-j}(s,t) B^j(s,t) \eta^k(s,t)}{C^k(s,t)} \\
&\qquad = \frac{1}{C^m(s,t)} \sum_{j=0}^{\min\{m,l\}} (-1)^{l-j} \binom{m}{j} \binom{-m}{l-j} \frac{A^{m-j}(s,t) B^j(s,t) \eta^{l-j}(s,t)}{C^{l-j}(s,t)} \\
&\qquad\qquad = \frac{A^m(s,t)\eta^l(s,t)}{C^{m+l}(s,t)} \sum_{j=0}^{\min\{m,l\}} (-1)^{l-j} \binom{m}{j} \binom{-m}{l-j} \left(\frac{B(s,t)C(s,t)}{A(s,t)\eta(s,t)}\right)^j.
\end{aligned}$$

Returning to our original notation,

$$\frac{1}{l!}\frac{\partial^l F_m}{\partial z^l}(1-f,s,t) = \frac{(q(s,t)+(1-f)(1-q(s,t)-\eta(s,t)))^m \eta^l(s,t)}{(1-(1-f)\eta(s,t))^{m+l}}$$
$$\times \sum_{j=0}^{\min\{m,l\}} (-1)^{l-j}\binom{m}{j}\binom{-m}{l-j}\left(\frac{(1-q(s,t)-\eta(s,t))(1-(1-f)\eta(s,t))}{(q(s,t)+(1-f)(1-q(s,t)-\eta(s,t)))\eta(s,t)}\right)^j$$
$$= \frac{(q(s,t)+(1-f)(1-q(s,t)-\eta(s,t)))^m \eta^l(s,t)}{(1-(1-f)\eta(s,t))^{m+l}}$$
$$\times \sum_{j=0}^{\min\{m,l\}} \binom{m}{j}\binom{m+l-j-1}{l-j}\left(\frac{(1-q(s,t)-\eta(s,t))(1-(1-f)\eta(s,t))}{(q(s,t)+(1-f)(1-q(s,t)-\eta(s,t)))\eta(s,t)}\right)^j$$

and

$$\frac{1}{l!}\frac{\partial^l F_m}{\partial z^l}(1-f,t,T_{\mathrm{mrca}})) = \frac{(q(t,T_{mcra})+(1-f)(1-q(t,T_{mcra})-\eta(t,T_{mcra})))^m \eta^l(t,T_{mcra})}{(1-(1-f)\eta(t,T_{mcra}))^{m+l}}$$
$$\times \sum_{j=0}^{\min\{m,l\}} (-1)^{l-j}\binom{m}{j}\binom{-m}{l-j}\left(\frac{(1-q(t,T_{mcra})-\eta(t,T_{mcra}))(1-(1-f)\eta(t,T_{mcra}))}{(q(t,T_{mcra})+(1-f)(1-q(t,T_{mcra})-\eta(t,T_{mcra})))\eta(t,T_{mcra})}\right)^j,$$

which gives us an expression for the derivatives.

### E Confidence Intervals

Immediately from the above, we have

$$\mathbb{P}(m_1 \le N(t) \le m_2 | N(s)=x, N_{\mathrm{obs}}(T_{\mathrm{mrca}})=l)$$
$$= \sum_{m=m_1}^{m_2} \mathbb{P}(N(t)=m|N(s)=x, N_{\mathrm{obs}}(T_{\mathrm{mrca}})=l)$$
$$= \sum_{m=m_1}^{m_2} \frac{\frac{\partial^l F_m}{\partial z^l}(1-f,t,T_{\mathrm{mrca}})}{\frac{\partial^l F_x}{\partial z^l}(1-f,s,T_{\mathrm{mrca}})}\mathbb{P}(N(t)=m|N(s)=x),$$

or, when $N(T_{\mathrm{mrca}})$ is known, as in (A.3) and (A.4), we have

$$\mathbb{P}(m_1 \le N(t) \le m_2 | N(s)=x, N(T_{\mathrm{mrca}})=n)$$
$$= \sum_{m=m_1}^{m_2} \frac{\mathbb{P}(N(T_{\mathrm{mrca}})=n|N(t)=m)}{\mathbb{P}(N(T_{\mathrm{mrca}})=n|N(s)=x)}\mathbb{P}(N(t)=m|N(s)=x).$$

These may be used to determine a minimal interval $m_1 \le m \le m_2$ such that

$$\mathbb{P}(m_1 \le N(t) \le m_2 | N(s)=x, N_{\mathrm{obs}}(T_{\mathrm{mrca}})=l) \ge 1-\varepsilon,$$

for any $\varepsilon > 0$. The values $m_1$ and $m_2$ need not be unique; one way to approach this systematically would be to choose the value of $m$ such that the probability that $N(t)=m$ is maximal, and then inductively generate the interval by subsequently including the natural number with maximum probability adjacent to the interval until the threshold $1-\varepsilon$ is attained. We could choose our initial $m$ by first determining the mean value of $N(t)$, which we discuss below. In the current paper, we did not follow this procedure, but rather computed the probabilities for all $m$, until their sum was indistinguishable from 1. We then retained the values of $m$ with highest probability that collectively summed up to 0.95.

### F Mean Number of Lines

We can also use the previous to obtain a closed form for the expected number of species at time $t$:

$$
\begin{aligned}
&\mathbb{E}\left[N(t)|N(s)=x, N_{\text{obs}}(T_{\text{mrca}})=l\right] \\
&\qquad = \sum_{m=1}^{\infty} m \frac{\frac{\partial^l F_m}{\partial z^l}(1-f,t,T_{\text{mrca}})}{\frac{\partial^l F_x}{\partial z^l}(1-f,s,T_{\text{mrca}})} \mathbb{P}\left(N(t)=m|N(s)=x\right) \\
&\qquad = \frac{\frac{\partial^l}{\partial z^l}\Big|_{z=1-f}\left(\sum_{m=1}^{\infty} m\mathbb{P}\left(N(t)=m|N(s)=x\right)F(z,t,T_{\text{mrca}})^m\right)}{\frac{\partial^l F_x}{\partial z^l}(1-f,s,T_{\text{mrca}})} \\
&\qquad = \frac{\frac{\partial^l}{\partial z^l}\Big|_{z=1-f}\left(F(z,t,T_{\text{mrca}})\frac{\partial F_x}{\partial z}\left(F(z,t,T_{\text{mrca}}),s,t\right)\right)}{\frac{\partial^l F_x}{\partial z^l}(1-f,s,T_{\text{mrca}})}.
\end{aligned}
$$

Now, observing that

$$F_x\left(F(z,t,T_{\text{mrca}}),s,t\right) = \left(F(F(z,t,T_{\text{mrca}}),s,t)\right)^x = \left(F(z,s,T_{\text{mrca}})\right)^x = F_x(z,s,T_{\text{mrca}}),$$

we have that

$$\frac{\partial F_x}{\partial z}\left(F(z,t,T_{\text{mrca}}),s,t\right)\frac{\partial F}{\partial z}(z,t,T_{\text{mrca}}) = \frac{\partial F_x}{\partial z}(z,s,T_{\text{mrca}}),$$

so the mean value reduces to

$$\frac{\frac{\partial^l}{\partial z^l}\Big|_{z=1-f}\left(\frac{F(z,t,T_{\text{mrca}})\frac{\partial F_x}{\partial z}(z,s,T_{\text{mrca}})}{\frac{\partial F}{\partial z}(z,t,T_{\text{mrca}})}\right)}{\frac{\partial^l F_x}{\partial z^l}(1-f,s,T_{\text{mrca}})} = \frac{\frac{\partial^l}{\partial z^l}\Big|_{z=1-f}\left(\frac{\partial \ln F}{\partial z}(z,t,T_{\text{mrca}})\frac{\partial F_x}{\partial z}(z,s,T_{\text{mrca}})\right)}{\frac{\partial^l F_x}{\partial z^l}(1-f,s,T_{\text{mrca}})},$$

or $\frac{\frac{\partial^n}{\partial z^n}\Big|_{z=0}\left(\frac{\partial \ln F}{\partial z}(z,t,T_{\text{mrca}})\frac{\partial F_x}{\partial z}(z,s,T_{\text{mrca}})\right)}{\frac{\partial^n F_x}{\partial z^n}(0,s,T_{\text{mrca}})}$ when one knows $N(T_{\text{mrca}})=n$.

In the current paper, we did not use these expressions, but rather computed the expected number of species at any time by computing $\sum_m m\mathbb{P}(N(t)=m)$ for all m values such that $\sum_m \mathbb{P}(N(t)=m)$ is at least 0.99.

### G Rates for the Conditioned Process

Finally, we observe that we may characterise the process conditioned on $N_{\text{obs}}(T_{\text{mcra}})$ and $N(s)$ as another time-inhomogeneous birth-and-death process, only now with frequency dependent rates.

Using the Markov property, we have

$$\begin{aligned}
&\mathbb{P}\left(N(t+h)=m+k|N(t)=m, N(s)=x, N_{\text{obs}}(T_{\text{mrca}})=l\right)\\
&\quad=\mathbb{P}\left(N(t+h)=m+k|N(t)=m, N_{\text{obs}}(T_{\text{mrca}})=l\right)\\
&\quad=\frac{\frac{\partial^l F_{m+k}}{\partial z^l}(1-f,t+h,T_{\text{mrca}})}{\frac{\partial^l F_m}{\partial z^l}(1-f,t,T_{\text{mrca}})}\mathbb{P}\left(N(t+h)=m+k|N(t)=m\right)
\end{aligned}$$

Now,

$$\mathbb{P}\left(N(t+h)=m+k|N(t)=m\right)=\begin{cases}\lambda(t)mh+o(h) & \text{if } k=1,\\ \mu(t)mh+o(h) & \text{if } k=-1,\\ 1-(\lambda(t)+\mu(t))mh+o(h) & \text{if } k=0, \text{ and,}\\ 0 & \text{otherwise}\end{cases}$$

whilst

$$\frac{\partial^l F_{m+k}}{\partial z^l}(1-f,t+h,T_{\text{mrca}})=\frac{\partial^l F_m}{\partial z^l}(1-f,t,T_{\text{mrca}})+\mathcal{O}(h),$$

so that

$$\begin{aligned}
&\mathbb{P}\left(N(t+h)=m+k|N(t)=m, N(s)=x, N_{\text{obs}}(T_{\text{mrca}})=l\right)\\
&=\begin{cases}\frac{\frac{\partial^l F_{m+1}}{\partial z^l}(1-f,t,T_{\text{mrca}})}{\frac{\partial^l F_m}{\partial z^l}(1-f,t,T_{\text{mrca}})}\lambda(t)mh+o(h) & \text{if } k=1,\\
\frac{\frac{\partial^l F_{m-1}}{\partial z^l}(1-f,t,T_{\text{mrca}})}{\frac{\partial^l F_m}{\partial z^l}(1-f,t,T_{\text{mrca}})}\mu(t)mh+o(h) & \text{if } k=-1,\\
\left(1-\frac{\frac{\partial^l F_{m+1}}{\partial z^l}(1-f,t,T_{\text{mrca}})}{\frac{\partial^l F_m}{\partial z^l}(1-f,t,T_{\text{mrca}})}\lambda(t)\right. & \\
\qquad\left.+\frac{\frac{\partial^l F_{m-1}}{\partial z^l}(1-f,t,T_{\text{mrca}})}{\frac{\partial^l F_m}{\partial z^l}(1-f,t,T_{\text{mrca}})}\mu(t)\right)mh+o(h) & \text{if } k=0, \text{ and}\\
0 & \text{otherwise.}\end{cases}
\end{aligned}$$

Similarly, using (A.3) and (A.4), to reduce to the case when $N(T_{\text{mrca}})$ is known, we have

$$\begin{aligned}
&\mathbb{P}\left(N(t+h)=m+k|N(t)=m, N(s)=x, N(T_{\text{mrca}})=n\right)\\
&=\begin{cases}\frac{\mathbb{P}(N(T_{\text{mrca}})=n|N(t)=m+1)}{\mathbb{P}(N(T_{\text{mrca}})=n|N(s)=m)}\lambda(t)mh+o(h) & \text{if } k=1,\\
\frac{\mathbb{P}(N(T_{\text{mrca}})=n|N(t)=m-1)}{\mathbb{P}(N(T_{\text{mrca}})=n|N(s)=m)}\mu(t)mh+o(h) & \text{if } k=-1,\\
1-\left(\frac{\mathbb{P}\left(N(T_{\text{mrca}})=n|N(t)=m+1\right)}{\mathbb{P}\left(N(T_{\text{mrca}})=n|N(s)=m\right)}\lambda(t)\right. & \\
\qquad\left.+\frac{\mathbb{P}\left(N(T_{\text{mrca}})=n|N(t)=m-1\right)}{\mathbb{P}\left(N(T_{\text{mrca}})=n|N(s)=m\right)}\mu(t)\right)mh+o(h) & \text{if } k=0, \text{ and}\\
0 & \text{otherwise.}\end{cases}
\end{aligned}$$

This gives us an efficient way of simulating specific realisations of the diversity through time curve, given the extinction and speciation rate functions, $T_{\mathrm{mrca}}$, and either the number of extant species at present or the number of observed species at present and $f$.
