## Supplementary Material for "Estimating Diversity Through Time using Molecular Phylogenies: Old and Species-Poor Frog Families are the Remnants of a Diverse Past"

Billaud, O., Moen, D., Parsons, T. L., Morlon, H.

### Supplementary Figures and Tables:

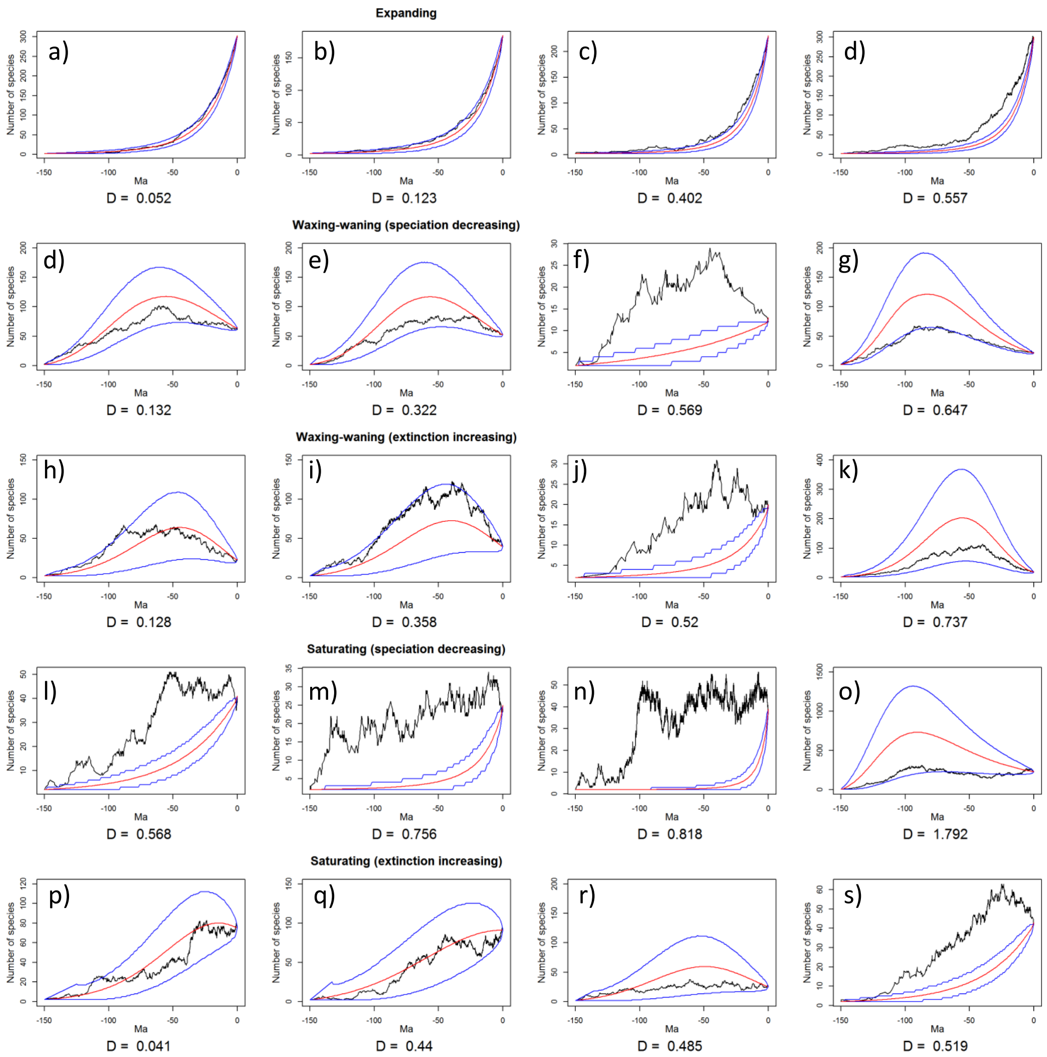


**Supplementary Figure 1** Illustration of simulated diversity trajectories, with estimated DTT curves (in red), confidence intervals (in blue), and associated global errors D for the five diversification scenarios considered in the paper. Most of the largest errors (D>0.5) arise from improper model selection (f, j, l, m, n, o, s), but they can also arise from biases in parameter estimation even when the proper model is selected (d, g, k). Cases with a global error below 0.5 in general corresponds to cases when the confidence interval encompasses the true diversity trajectory. Confidence intervals are sometimes so wide that they encompass the true diversity trajectory even if the global error is large (k).

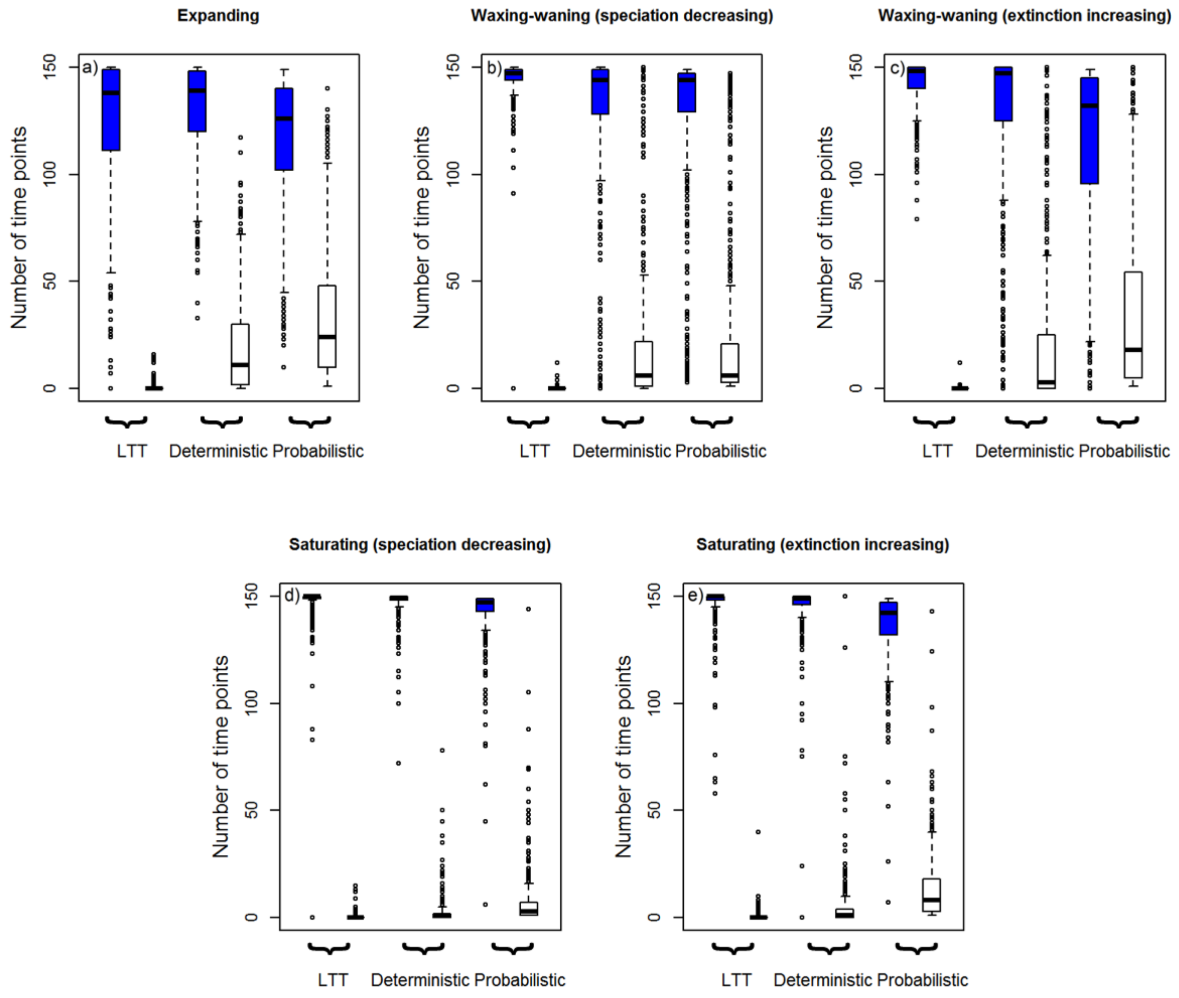


**Supplementary Figure 2** Number of over- (in white) versus under-estimations (in blue) across the 150 time points for which estimates were computed, for trees simulated under the five diversification scenarios considered in the paper, and when using each of the three diversity-through-time estimates. Boxplots represent the median, $1^{st}$ and $4^{th}$ quartile over 400 simulations, whiskers represent the lowest (and highest) datum still within 1.5 interquantile range of the lower (resp.upper) quartile, and dots represent outliers.

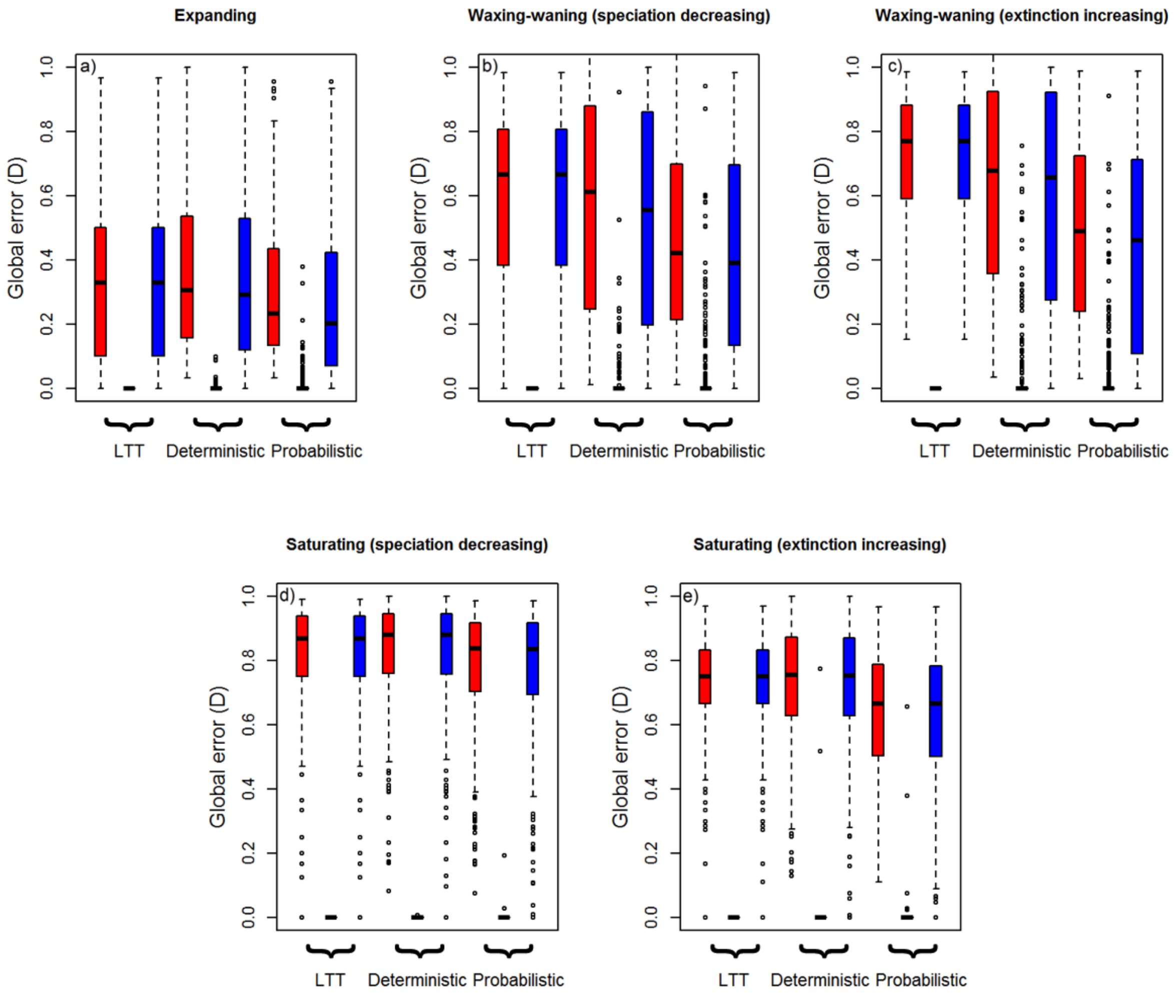


**Supplementary Figure 3** Global error D (in red), overestimation $D^+$ (in white) and underestimation $D^-$ (in blue) for trees simulated under the five diversification scenarios considered in the paper when using each of the three diversity-through-time estimates. Boxplots represent the median, $1^{st}$ and $4^{th}$ quartile over 400 simulations, whiskers represent the lowest (and highest) datum still within 1.5 interquantile range of the lower (resp.upper) quartile, and dots represent outliers.

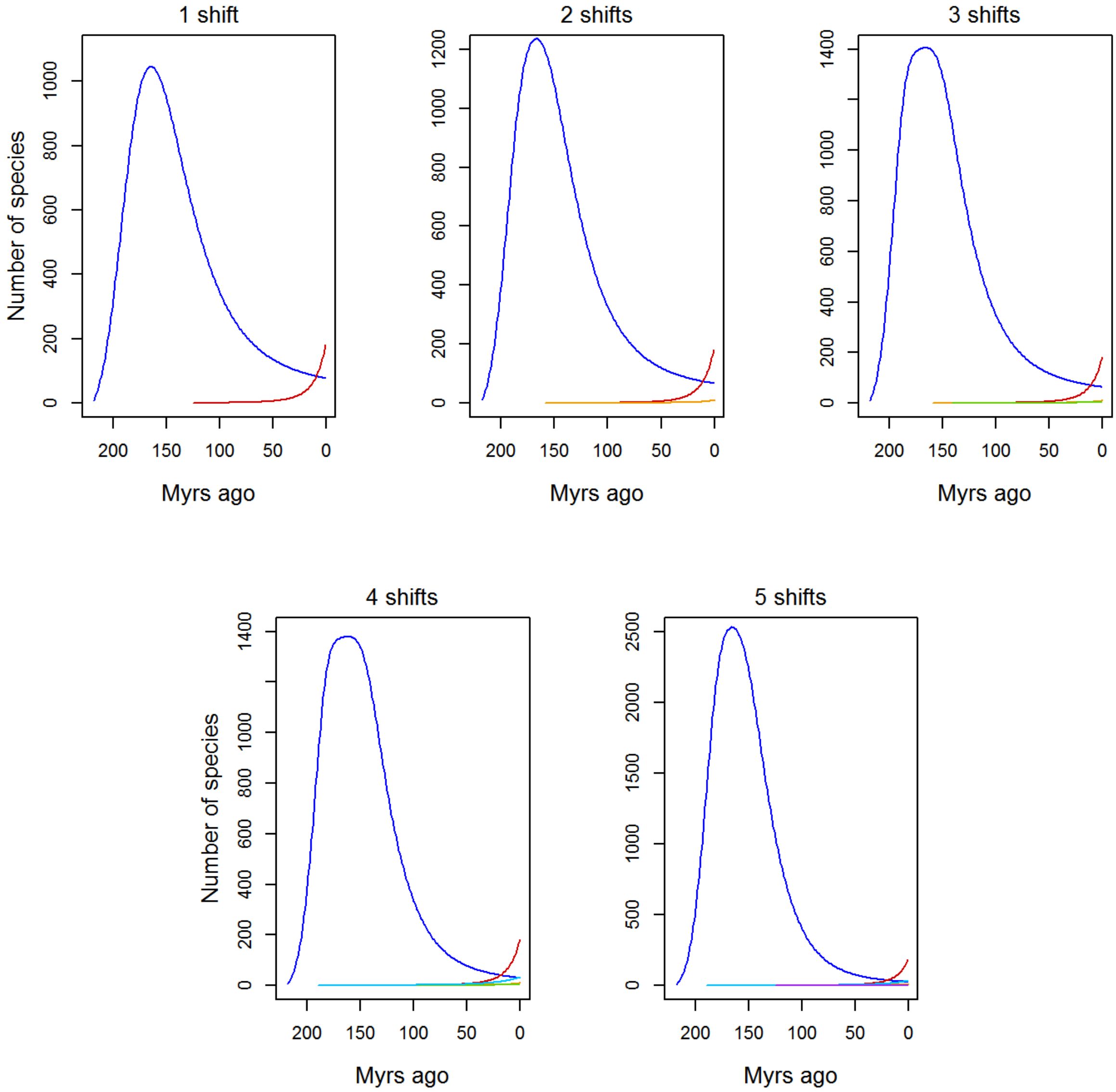


**Supplementary Figure 4** Inferred diversity trajectories of archaeobatrachian frogs for an increasing number of assumed rate shifts. The blue curve depicts the expected diversity-through-time curve corresponding to the 'backbone' phylogeny (i.e. the phylogeny that excludes the family(ies) for which a diversification rate shift has been inferred). The colored curves depict the expected diversity-through-time curves corresponding to the families subtending diversification rate shifts (red: Megophryidae; orange: Bombinatoridae; green: Pelodytidae; blue: Pipidae; purple: Pelobatidae)

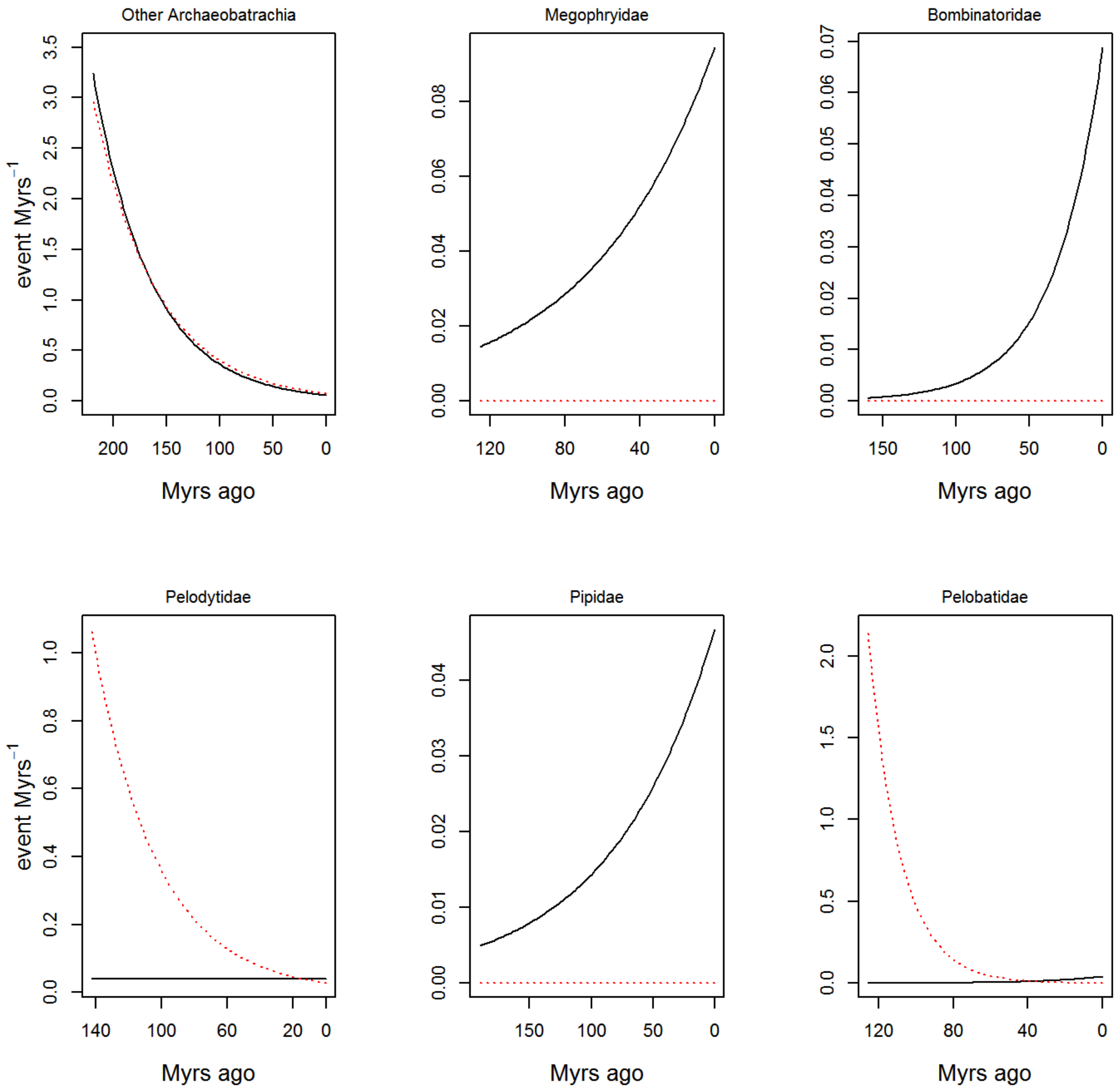


**Supplementary Figure 5** Inferred rates of speciation (in black) and extinction (dashed red) through time for the "backbone" Archaeobatrachia phylogeny and the five families subtending diversification rate shifts.

**Supplementary Table 1**

| Number of shifts | Family concerned by the new shift | P-value |
|---|---|---|
| 1 | Megophryidae | **4.91E-08** |
| 2 | Bombinatoridae | **2.34e-03** |
| 3 | Pelodytidae | **7.57e-0.3** |
| 4 | Pipidae | **1.43e-03** |
| 5 | Pelobatidae | **9.78e-03** |
| 6 | Leiopelmatidae | 5.03e-02 |

**Support for successive diversification rate shifts in the Archaeobatrachia phylogeny.** P-values represent *P*-values of the likelihood ratio test between the corresponding model and the simpler model with one less shift. Values in bold are significant
